# Food web complexity modulates environmental impacts on food chain length

**DOI:** 10.1101/2023.04.29.538714

**Authors:** Shota Shibasaki, Akira Terui

**Affiliations:** Department of Biology, University of North Carolina at Greensboro, USA

**Keywords:** food web, motif, network analysis, biodiversity, mathematical modeling, trophic interactions

## Abstract

Food chain length (FCL), a crucial aspect of biodiversity, has long been debated for its determinants. Previous studies proposed resource availability, disturbance, and ecosystem size as environmental drivers. However, studies using stable isotope approaches have shown inconsistent results, indicating missing links between environmental drivers and FCL. Here, we hypothesized that species richness and motifs (i.e., 3-species subgraphs) modulated environmental effects on FCL. Combining empirical food webs with our *N* -species food web model, we found that FCL disproportionately changed at low species richness with saturation at high species richness. This functional response was essential to the interdependent effects of disturbance and ecosystem size in our model. Disturbance more strongly regulated FCL in smaller ecosystems, where species richness was low. Similarly, increasing ecosystem size enhanced FCL under strong, but not weak, disturbance regimes. Our study suggests that internal food web structure should deepen our understanding of how FCL changes over environments.

## 1 Introduction

Food webs have been of ecologist’s interest for at least one hundred years (Elton, 1927). The food chain length (FCL) is a crucial aspect of food webs because FCL represents a vertical dimension of biodiversity (Wang and Brose, 2018) and is associated with top-down effects, primary productions, and toxin contamination in predators (Kidd et al., 1995; Pace et al., 1999; Wang and Brose, 2018). Three environmental factors are proposed as drivers of FCL: resource availability (Oksanen et al., 1981), disturbances (Pimm and Lawton, 1977), and ecosystem sizes (Post et al., 2000). Previous studies show, however, inconsistent results on whether these three drivers change FCL (Briand and Cohen, 1987; Jake et al., 2007; Takimoto et al., 2008; Doi et al., 2009; McHugh et al., 2010; Sabo et al., 2009, 2010; Takimoto and Post, 2013), resulting in a long-lasting debate on the environmental determinants of FCL.

The ongoing debate regarding context dependency in FCL suggests the existence of missing modulators between environmental drivers and FCL. This potential shortcoming may be tied to the current common approach in FCL research, i.e., stable isotopes. Traditionally, empirical FCLs were assessed by measuring the feeding links connecting the basal and apex species within a food web (see Briand and Cohen (1987) for example); however, this method (so-called “connectance food web”) is sensitive to issues arising from uneven and biased taxonomic resolutions within the food web nodes (Winemiller, 2007; Pringle and Hutchinson, 2020). The stable isotope approach elegantly addressed this problem by integrating food web interactions (Layman et al., 2012), whose signatures are condensed into “top” (i.e., top predator) and “bottom” (primary producers or consumers) of the food web as stable isotope ratios. The isotopic difference between the two approximates the vertical structure of the food web and the energy flow therein. Importantly, it naturally accounts for the inherent complexity of predator-prey interactions, such as omnivory. These distinctive features rendered stable isotopes a promising tool to quantify FCL in natural systems (Post et al., 2000; Post, 2002; Jake et al., 2007; Layman et al., 2007; Takimoto et al., 2008; Sabo et al., 2010; Sullivan et al., 2015).

While we acknowledge the major advantage of stable isotopes in FCL research, however, it lumps the internal food web structure that may lead to the emergent relationships between FCL and environmental factors (Fig. 1). In addition, existing experiments and theoretical frameworks often oversimplify natural food webs, typically with three- to four-species communities (Pimm and Lawton, 1977; Hastings, 1979; Diehl and Feissel, 2001; Takimoto et al., 2012; Ward and McCann, 2017; Doi and Hillebrand, 2019), or assuming only chain networks (Liao et al., 2016; Jonsson, 2017; Terui and Nishijima, 2019; Wang et al., 2021). Although there are a few noteworthy exceptions (Martinez and Lawton, 1995; Neutel et al., 2007), it remains unclear how the internal structure of food webs alters the relationship between FCL and external environmental drivers. Here, we employ the classical “connectance food web” approach to shed light on the role of food web structure in shaping the relationship between FCL and external environmental drivers.

**Figure 1:**
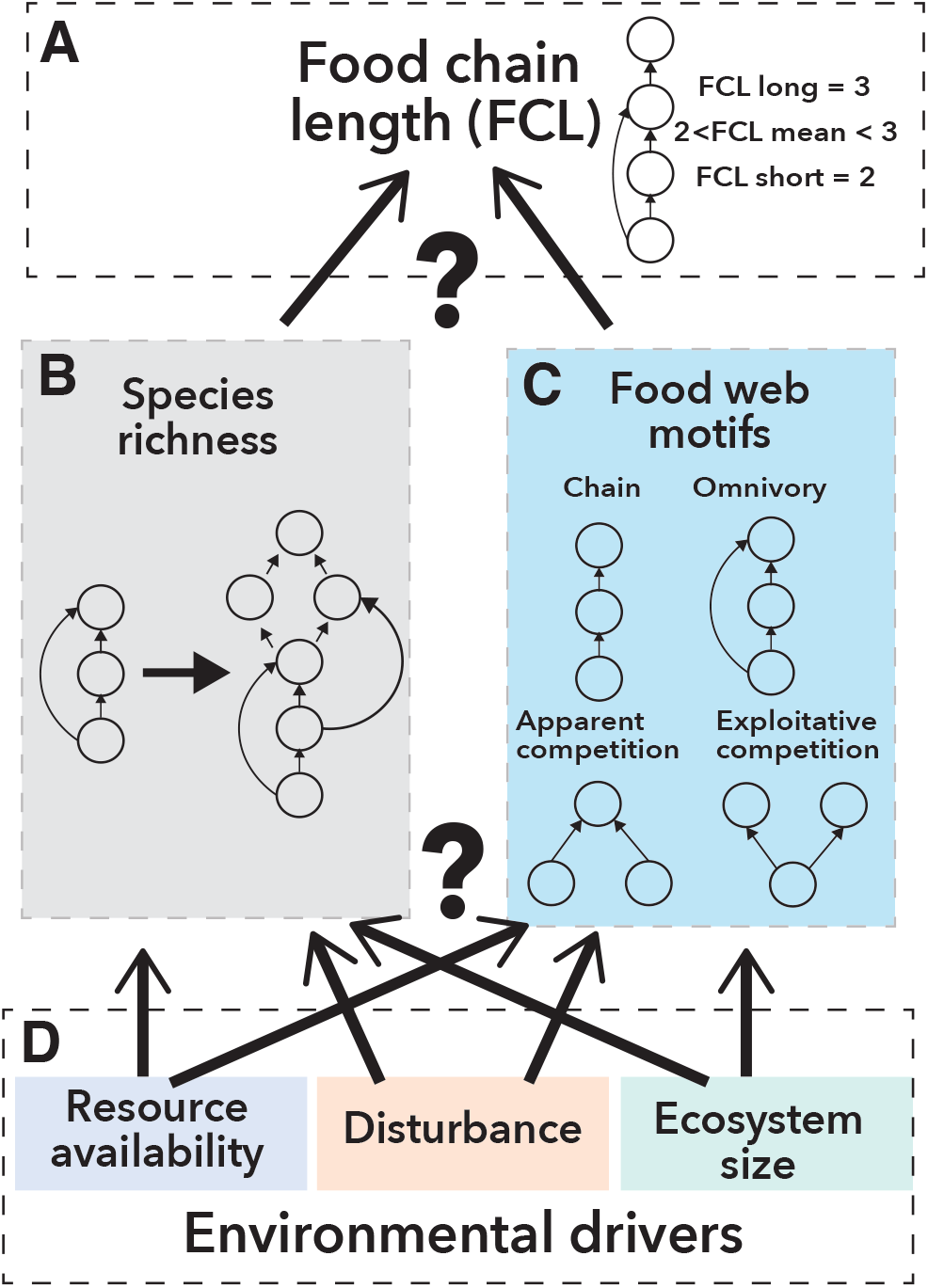
Schematic representations of research questions. We investigate whether and how species richness (panel B) and food we motifs (panel C; three-species chain, omnivory, apparent competition, and exploitative competition) modulate the relationship between the environmental drivers (panel D; resource availability, disturbance, and ecosystem size) and FCL, (panel A). FCL is defined in three ways depending on how trophic positions are defined. If trophic levels are defined by the one plus the longest-path length from basal species (FCL long), FCL of the food web on the top panel is three. If trophic positions are given by one plus the mean of the prey species’ trophic positions (FCL mean), its FCL is between two and three. FCL short measures trophic positions as one plus the shortest-path length from basal species; FCL short of the top food web is two. In the main text, we focused on FCL mean. See SI 2 in cases of the rest FCL definitions.

A parallel line of research has documented how species richness and motifs, i.e., three-species subnetworks within a food web (Milo et al., 2002), alter the stability of food webs, which should be linked to the maintenance of long food chains. As such, a reasonable first step is to explore the processes through which these internal factors mediate the impacts of environmental controls on FCL (Figs. 1 B and C). Increasing species richness enhances FCL if an inserted species either occupies the higher trophic positions than resident species or increases trophic position(s) of resident top predator(s) (Post and Takimoto, 2007) (but see SI 6 for the analysis of how “species richness” increases FCL in the random graph); however, decades of “diversity-stability” debates (McCann, 2000) revealed that random species assembly destabilizes food webs (Gardner and Ashby, 1970; May, 1972; Allesina and Tang, 2012), and the non-randomness of assembly rules is the critical property of persistent food webs. Previous studies show that the following four motifs are dominant in empirical food webs (Stouffer et al., 2007; Borrelli, 2015; Monteiro and Faria, 2016) and relate to the persistence of food webs (Cirtwill and Wootton, 2022; Stouffer and Bascompte, 2010; Monteiro and Faria, 2016): three-species chain, omnivory, apparent competition, and exploitative (or direct) competition (Fig. 1C). Notably, these motifs differ in their FCL; the chain motif is longer than the two competition motifs, and the length of the omnivory motif is between the chain and the competition.

In this study, we investigate whether and how species richness (or technically “operational species richness” defined in Section 2.1) and food web motifs modulate the relationship between FCL and the three environmental drivers (i.e., resource availability, disturbances, and ecosystem sizes). We begin with the analysis of an empirical food web database (Cohen, 2010) to assess the association between FCL and internal food web properties (operational species richness and food web motifs). With simulations of species-rich food webs, we further elaborated on how the association alters the relationship between the three environmental drivers and FCL. Our analysis suggests that species richness modulates environmental effects on FCL because of a nonlinear relationship between species richness and FCL. This may explain the inconsistent patterns between the environmental drivers and FCL in previous studies.

## 2 Methods

### 2.1 Definitions and terminologies

In this study, we refer to the number of nodes in a food web as “operational species richness” because the number of nodes does not always correspond to the number of biological species (species richness). In empirical food webs, each node may include multiple biological species that share trophic status (i.e., trophic species) (Pringle and Hutchinson, 2020) or are taxonomically close due to the limited resolutions; the issue of taxonomic resolution in food webs has been discussed in previous studies (Sugihara et al., 1997; Yodzis and Winemiller, 1999). Alternatively, a single species can be subdivided into multiple nodes if the species has size-or stage-dependent trophic interactions. Individuals of such species should be assigned to different nodes to reflect their variations in diet (Cohen et al., 1993; Huxham et al., 1995); otherwise, information in a food web is lost (Luczkovich et al., 2003). The nodes in our simulations do not necessarily represent the biological species for the same reasons; our model does not include the traits that define either species’ boundaries (e.g., reproductive isolation) or size-dependent trophic interactions. We, therefore, use the term operational species richness in this manuscript. Similarly, “species” in this manuscript refers to a unit of entity (i.e., node) in a food web.

We defined FCL as the maximum trophic position within a food web minus one (Post, 2002). However, multiple definitions of trophic positions have been used in the literature. We used three of those defined in connectance (or binary) food webs: *mean, short*, and *long* trophic positions (hereafter, FCL *mean, short*, and *long* for corresponding FCLs). The *mean* trophic position is given by the mean trophic position of prey species plus one, while the *short* and *long* trophic positions are defined as the shortest- and longest-path length from the basal species plus one, respectively. These definitions can lead to different FCLs of the identical food web (see Fig. 1 A as an example). We used these three definitions to analyze factors that consistently affect FCL. The main text, however, focuses on the results of FCL mean. See SI 2 for the results with the two other definitions. In FCL mean, a trophic position of a predator species can reflect how much each prey species is eaten by the focal predator, although such information is not always available in empirical data. The other two definitions of FCL require only binary food webs, which contain the presence and absence of prey-predator interactions among species. FCL short can be measured in all food webs while FCL long can be used only in acyclic food webs (e.g., food webs without cannibalism and mutual predations). See also SI 1.1 for further details on FCL measures.

### 2.2 Empirical database

We analyzed 213 empirical food webs (from WEB1 to WEB213) in the database of Cohen (2010). This database contains adjacency matrices of food webs across many ecosystem classes (Fig. S18). This enables us to calculate FCL, species richness, and the proportion of different three-species motifs in empirical food webs. Our analysis focuses on the following three-species motifs: chain, omnivory, apparent competition, and exploitative competition. We focused on these motifs because they were dominant in the empirical food webs as reported in a previous study (Camacho et al., 2007). In addition, this result implies that most empirical food webs (203 out of 213) in this study are acyclic, where FCL long can be measured. See also SI 1.2 for more details.

common issueS in the database of Cohen (2010) are (1) the low taxonomic resolution and (2) binary prey-predator interactions. To address these typical issues in the food web database, we analyzed additional 11 food webs obtained from another database in SI 3, where higher taxonomic resolution and quantitative pre-predator information are available.

### 2.3 Mathematical model

#### 2.3.1 Generating food webs

While the empirical database provides the important links among FCL, species richness, and motifs, it does not contain data for resource availability, disturbances, or ecosystem sizes. This limited our ability to understand how environmental factors control FCL through the internal structures of food webs. To complement this limitation, we simulated the community dynamics of an *N* -species food web model that enables us to analyze how the environmental drivers affect FCL through species richness and food web motifs. In previous studies, multiple algorithms are proposed to produce *N* -species food webs that mimic empirical food webs (Allesina et al., 2008; Caldarelli et al., 1998; Cattin et al., 2004; Cohen and Newman, 1985; Kondoh and Ninomiya, 2009; Rossberg et al., 2005, 2006; Stouffer et al., 2005, 2006; Williams and Martinez, 2000). We used the modified preferential prey model (Johnson et al., 2014) for three reasons. First, this algorithm generates only acyclic food webs. As explained in Section 2.1, we can use only acyclic food webs when we measure FCL long. Although the original model in Johnson et al. (2014) allows cannibalism, we modified the model to preclude cannibalism from our simulated food webs. Second, this model explicitly pre-determines the number of basal species in a food web. Although (generalized) cascade models (Cohen and Newman, 1985; Stouffer et al., 2005) also generate acyclic food webs, we cannot fix the number of basal species in these models. The number of basal species may change FCL short and long because trophic positions are measured by the path length from basal species. Finally, the modified preferential prey model generates food webs given the definition of trophic positions (see below). A similar model proposed by Kondoh and Ninomiya (2009) satisfies the conditions of acyclic models and the fixed number of basal species; however, this model cannot incorporate the difference in the definitions of trophic positions. We used the preferential prey model to generate food webs for these reasons.

In the preferential prey model, we first need to determine the maximum number of species *N*, the maximum number of basal species *B*, and the expected number of trophic links *l*. In this study, we set (*N, B, l*) = (32, 4, 113) with the following rationale. *N* = 32 is sufficiently large to generate variations in realized operational species richness (see Results), yet computational costs of stochastic simulations (see below) are reasonably small. *B* = 4 was used to avoid the entire loss of the basal species, which causes the global extinction of the food web. In addition, excessive numbers of basal species limit the observable variation in FCL because fewer species are expected to attain high trophic positions. In such cases, statistical analyses would be difficult. *l* = 113 was chosen to meet the connectance *l/N* ^2^ ≈0.11, following statistics in Dunne et al. (2002).

Second, after setting parameters of (*N, B, l*), species *i* = 1,…, *B* are assigned as basal species with a trophic position of one. For each non-basal species *i* = *B* + 1,…, *N*, its prey species are assigned as follows. Non-basal species *i* randomly chooses its first prey species from *j* = 1,… *B*, …, *i-*1. If species *i* has multiple prey species, additional prey species are stochastically assigned. Non-basal species *i* tends to choose additional prey species whose trophic positions are close to species *i*’s initial prey (species *j*), but this tendency is tuned by parameter *T* (*T→*0 represents species *i* consumes only species whose trophic positions are identical to species *j*, while *T→ ∞* represents species *i* choose prey species regardless of prey’s trophic positions). We set *T* = 1 in the main text. In the original preferential prey model, species’ trophic positions are determined by the mean of preys’ trophic positions plus one. We extended this model to define species’ trophic positions using the longest-or shortest-path length from the basal species. We sampled 30 food webs in each of the three definitions of the trophic positions to account for the stochasticity of the preferential prey model.

#### 2.3.2 Stochastic simulations of the ecological dynamics

Although the preferential prey model generates *N* -species food webs, the coexistence of *N* -species is not guaranteed. The model provides initial static food webs, which are subsequently “pruned” into realized food webs of coexisting species (i.e., subgraphs of the *N* -species food web) considering ecological dynamics. Hereafter, species richness and fractions of food web motifs in our simulations refer to those in realized food webs, whose values vary greatly among simulation replicates. To obtain the realized food webs, we simulated the ecological dynamics by extending the theory of island biogeography (MacArthur and Wilson, 1967). In the original theory of island biography, species colonization and extinction rates in a patch are constant. Gravel et al. (2011) extends this theory so that the colonization and extinction rates depend on the presence of prey and predator species in the patch, respectively. Saravia et al. (2022) showed that this modeling framework could reproduce the frequency of motifs and trophic positions that are statistically indistinguishable from empirical food webs. From this result, we assume that the extended theory of island biogeography model does not bias FCL and the food web motifs in realized food webs.

Our rationale for employing the theoretical framework of island biogeography is to make simulated results readily comparable to empirical food webs, which typically contain only the presence/absence information of constituent species. While previous studies implement population dynamics (Otto et al., 2007; Kondoh and Ninomiya, 2009), implementing population dynamics did not alter the results in the main text (See SI 4).

In our model, we consider the presence (*P*_*i*_ = 1) and absence (*P*_*i*_ = 0) of species *i* in a single patch (i.e., “island”) with species migration from the external permanent species pool (i.e., “mainland”). The colonization-extinction process is defined by the following reactions:

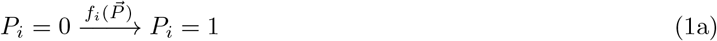

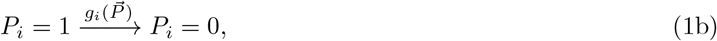

where *f*_*i*_ is the colonization rate and *g*_*i*_ is the extinction rate. These rates depend on the presence of other species 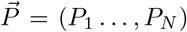 and environments. If species *i* is a basal species (1 ≤*i≤B*), the colonization rate depends on the resource availability, the ecosystem size, and the presence of other basal species:

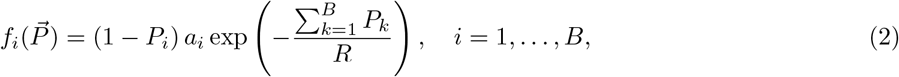

where *a*_*i*_ is the maximum colonization rate of basal species *i* in the absence of other basal species (i.e, no competitors for resource), and *R* quantifies the resource availability in the patch. The resource availability *R* defines how rapidly the colonization rate decreases as the number of other basal species increases.

For non-basal species *i* (*B* + 1≤ *i* ≤ *N*), on the other hand, the colonization rate depends on the presence of prey species:

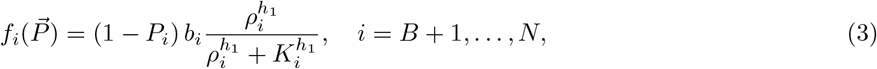

where *b*_*i*_ is the maximum colonization rate of non-basal species *i, ρ*_*i*_ is the number of species *i*’s prey species in the patch, *K* gives the number of prey species that defines the half-max colonization rate, and *h*_1_ is the Hill coefficient that determines the shape of the function. The maximum colonization rates (*a*_*i*_ and *b*_*i*_) reflect the ecosystem size because species would migrate more frequently into a larger patch than a smaller one (i.e., the target effect). In SI 5, we implemented alternative ecosystem size’s effect as decreasing the extinction rate due to predation *c*_*i*_ (see below), instead of increasing colonization rates (*a*_*i*_ and *b*_*i*_). This idea can be justified when population abundances are fixed regardless of the system size. In such cases, population densities decrease over ecosystem size, resulting in lower encounter rates between prey and predatorsn(Ward and McCann, 2017). SI 5 shows qualitatively similar results to the main text with this implementation.

The extinction rate is composed of predation, lack of prey, and disturbances (*e*_*i*_):

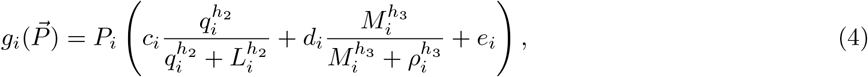

where *c*_*i*_ and *d*_*i*_ represent maximum extinction rates of species *i* due to abundant predators and scarce prey, respectively, *q*_*i*_ is the number of species *i*’s predator in the patch, *L*_*i*_ gives the number of predator species that defines the half-max extinction rate due to the predation pressure, *M*_*i*_ gives the number of prey species that defines the half-max extinction rate due to the lack of prey, and *h*_2_ and *h*_3_ are the Hill coefficients. For top predators, extinction due to predators never occurs (*c*_*i*_ = 0), while the basal species has *d*_*i*_ = 0 as they do not have any prey species. The default values and the descriptions of the parameters are listed in Table 1. In the main text, we focused on the effect of two environmental drivers: disturbance and ecosystem size. This is because our preliminary results (SI 7) suggest that resource availability showed no correlations. Although species would differ in their parameter values (e.g., colonization rates *a*_*i*_ and *b*_*i*_, sensitivity to disturbance *e*_*i*_, etc) in nature, we fixed the parameter values across the species to focus on the effects of network structures of food webs. We replicated the simulations 20 times for each combination of a food web and a parameter set. In total, we analyzed 30 food webs ×20 replicates ×72 parameter sets (variations in *a*_*i*_ and *e*_*i*_, see Table 1)= 43200 data in each definition of FCL.

**Table 1:**
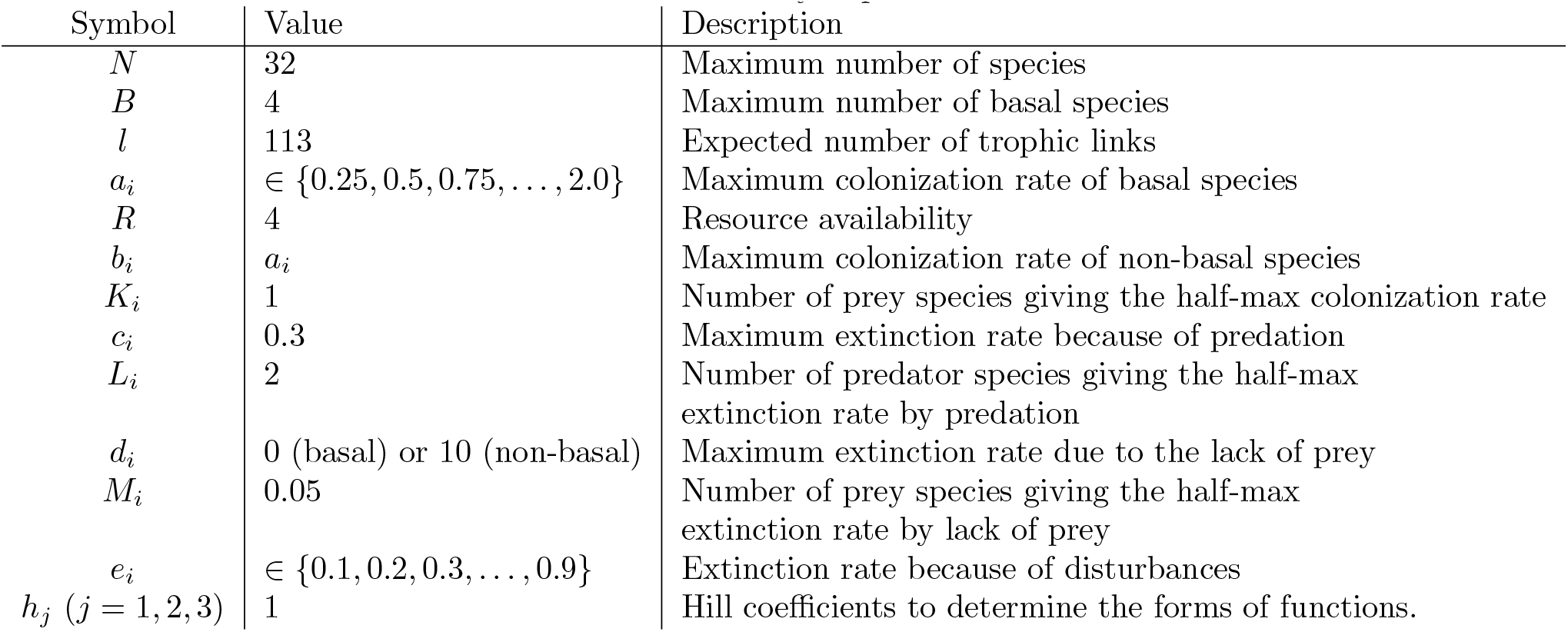
Summary of parameters.

The stochastic dynamics were implemented by Gillespie algorithm using GillespieSSA library (Pineda-Krch, 2008) version 0.6.2 in R version 4.2.1 (R Core Team, 2022). For each run of the simulations, all species are absent in the patch at the beginning: 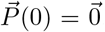. We continued the simulations until time 1, 000. The master equation of the system describes the dynamics of the probability distribution of 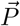 as follows:

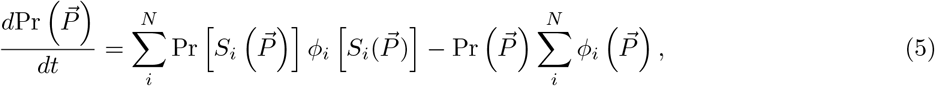

where Pr 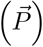 represents the probability density of presence-absence vector 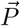, *S*_*i*_ is an operator to switch the presence of species *i* to the absence and vice versa without changing other species’ presence-absence information, i.e., 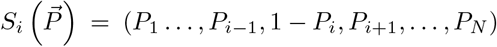, and 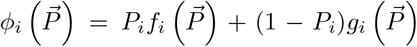 represents the switching rate of species *i* from presence to absence and vice versa. Intuitively, the first term of Eq (5) represents the rate that the species composition becomes 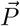 and the second term indicates the rate that species composition changes from 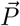.

After removing species with no prey from the realized food webs at the end of the simulations, FCL was measured as SI 1.1 explains, and the fractions of 13 motifs were calculated using igraph library (Csardi and Nepusz, 2006) version 1.3.5 in R. If all species go extinct, we regard FCL as -1 because FCL is zero when only basal species exist.

### 2.4 Sensitivity analysis

While we fixed parameter values of (*N, B, l, T*) in the main text, these parameter values affect the structure of entire food webs. To ensure the robustness of the results in the main text, we ran additional 7200 simulations in which parameters were randomly sampled from the following sets with equal probability: *N* ∈*{*16, 24, 32, 40, 48*},B∈{*2, 4, 6, 8*}, l/N* ^2^ ∈*{*0.9, 0.1, 0.11, 0.12, 0.13*}*, and *T ∈ {*0.1, 0.5, 1, 2, 10*}*. We varied the maximum operational species richness *N* within the range that the computational cost is not too huge. The range of the maximum number of basal species was chosen so that at least half of the species in a food web are non-basal species; otherwise, FCL would remain short. The range of connectance *l/N* was based on Dunne et al. (2002); the mean and standard deviation of connectance in empirical food webs are 0.11 and 0.09, respectively. The range of *T* was determined to generate a large variation of the entire food web structure following the results in Johnson et al. (2014). In this sensitivity analysis, we also randomly sampled the frequency of disturbance and ecosystem size while fixing resource availability and other parameter values as shown in Table 1. These simulations allow us to validate (i) whether operational species richness and the fraction of chain motifs correlate with FCL, and (ii) whether operational species richness modulates how environmental drivers affect FCL in diverse food web structures. The sensitivity analysis was performed in all three definitions of FCL.

### 2.5 Statistical analysis

#### 2.5.1 Statistical analysis on empirical data

We used regression analysis to examine how species richness and the fraction of the 3-species chain motif affects long, short, and mean FCLs (see Results). We assumed these two variables have either of the following three functional forms: linear, saturating, or quadratic. These functional forms were determined from the scatter plots against FCL (see Figs. S1 and S2). We generated fifteen nonlinear models for each definition of FCL: species richness or the fraction of the chain motif, or both affect FCL in either of the three functional forms (i.e., 2 × 3 + 3 × 3 = 15 models), respectively. The three functional forms are written as below:

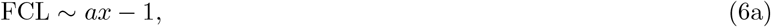

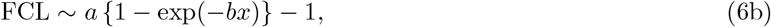

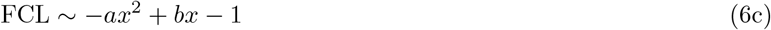

where *x* is either operational species richness or the fraction of the chain motifs in a food web, and coefficients *a* and *b* are assumed positive because of positive correlations between FCL and operational species richness or the fraction of chain motifs (see Figs. S1 and S2). The constant terms of the three models are fixed as *-*1 given our definition of FCL (see Subsection 2.3.2). In the saturating functional form (Eq (6b)), parameter *a* controls how much *x* (i.e., either species richness or the fraction of chain motifs) increases FCL because FCL converges to *a* − 1 in the limit of *x* → ∞ *1*. In the quadratic function (Eq (6c)), the maximum increase of FCL by *x* is either *b*^2^*/*(4*a*) (if *b<* 2*a*) or − *a* + *b* (otherwise).

We used nls() function with algorithm “port” in R to fit these models to the data from the empirical database. The lower boundaries of the coefficients were set zeros, while the upper boundaries were either maximum FCL in the data plus one (for *a* in the saturating function) or 10 (for the other coefficients). We chose the best models by calculating Akaike Information Criterion (AIC) of the regression models using AIC() function in R in each data. See Tables S6 - S8 for the ΔAIC and Table S3 for the best models and the coefficient values.

#### 2.5.2 Statistical analysis on simulation data

We repeated a similar regression analysis for our simulation data. We grouped simulated data according to the 30 food web structures, which were used as random effects in non-linear mixed effect models. We included the random effects in all coefficients in Eqs (6a) –(6c) because FCLs in different food webs may respond differently to changes in operational species richness and the fraction of chain motifs. We fitted the nonlinear mixed effect models to the data with nlme() function in nlme library (Pinheiro et al., 2023) version 3.1-162 in R. As mentioned above, the best model was chosen by AIC (see Tables S9 - S11 for ΔAIC).

The linear quantile mixed models (Tables S4 and S5) were used to quantify how disturbance and ecosystem size affect FCL and species richness, using lqmm() function in lqmm library (Geraci and Bottai, 2014) version 1.5.8 in R. We implemented the random effects of 30 entire food web structures on the slopes and the coefficients of the models. This analysis was intended to see whether the effects of disturbance on FCL differed depending on ecosystem size, and vice versa. Wilcoxon signed-rank test was also used to compare the logarithm of slopes of the linear quantile mixed models over disturbance or ecosystem size.

## 3 Results

### 3.1 Operational species richness non-linearly increases FCL

We first investigated whether operational species richness and the fractions of different food web motifs correlated with FCL to identify candidates for internal modulators. Operational species richness and three-species chain motifs positively correlated with FCL in all three definitions (Figs. S1A-F). The two competition motifs (i.e., apparent competition and exploitative competition), on the other hand, were negatively or non-significantly correlated with FCL (Figs. S1J-O). The correlations between FCL and the omnivory motif depended on the definition of trophic positions. Although omnivory increased FCL long (Figs. S1G), the correlation was weaker or non-significant with FCL mean (Figs. S1H) or short (Figs. S1I). These patterns were replicated in our simulated food webs (Fig. S2). The correlative analysis suggests that operational species richness and the chain motif may play key roles in modulating the environment–FCL relationships. We proceeded the further analyses to see how they relate to FCL and the potential environmental drivers.

The functional forms between FCL and internal food web structures (operational species richness and motifs) determine how environments dictate FCL. From Figs. S1 and S2, we compared three functional forms: linear, saturating, and quadratic functions. The best models (Table S3, see also Figs. S3 and S5 for FCL long and short, respectively) described the eect of operational species richness in the saturating functions both in empirical data and our simulations (Figs. 2A and B). This indicates that FCL changed disproportionately at low operational species richness with little changes at high operational species richness. This pattern did not change due to the difference in the entire food web structure; i.e., each 32-species food in simulations shows similar patterns over realized operational species richness (see dashed lines on Fig. 2B). The saturating functional form may be maintained in broad networks; it appeared in baseline models without species assembly (SI 6). In the meantime, quadratic models were selected as the best models for describing the functional relationships between FCL (except for FCL short, see Table S3) and the fraction of the chain motifs (Figs. 2C and D).

**Figure 2:**
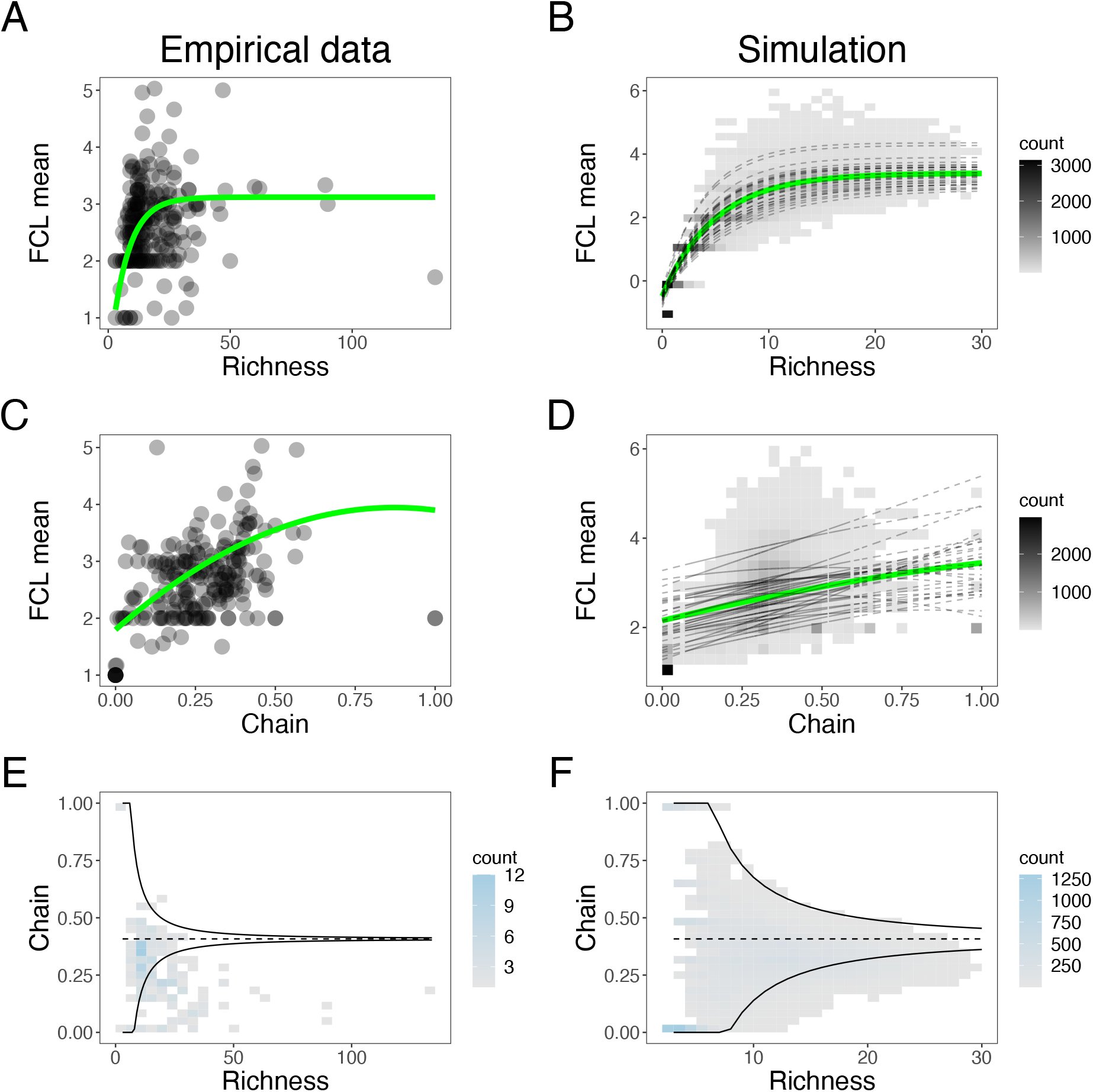
Species richness and the chain motifs relate to FCL. Relationships among operational species richness, the fraction of the three-species chain motifs, and FCL mean are shown (left: empirical data, right: our simulation data). A and B: operational species richness had saturating effects on FCL. The green curves represent the best models (Table S3) while fixing the fraction of the chain motif as the mean value. C and D: the fraction of the chain motif had quadratic effects on FCL. Similarly, the green curves represent the best models (Table S3) while fixing operational species richness as its mean value. Notably, the green curves on panels B and D represent only the fixed effects, and the black dashed lines include random effects. E and F: The fractions of the chain motifs over operational species richness are shown. The dashed line represents the mean fraction of the chain motifs in the random graph, and the solid curves represent mean *±* 2× ESD in the random graph. See SI 6 for more mathematical details.

The coefficients of best models (Table S3) indicated that operational species richness had larger maximum effects on FCL than chain motif; i.e., the coefficient *a* in Eq (6b) for operational species richness ranged from 2.85 (empirical FCL short) to 6.66 (simulated FCL long) while the maximum increase in FCL due to the chain motifs ranged from 1.23 (simulated FCL short) to 2.15 (empirical FCL long). However, species richness little increased FCL at high species richness because of the saturating function. If the fraction of chain motifs independently changes in operational species-rich food webs, the chain motifs may disproportionately influence FCL. To explore this possibility, we delved into how the fractions of the food web motifs changed over species richness. At high operational species richness, the fraction of chain motif was lower than expected in random graphs (the expected value *±*2 × expected standard deviation (ESD) in the random graphs) with no ecological rules in species assembly (Figs. 2E and F). In the random graph, the expected fractions of the motifs were constant but their ESD decreased over operational species richness (see SI 6 for mathematical details). These results imply that some regulation mechanisms hindered the persistence of chain motifs in species-rich food webs, limiting the motif effect on FCL. Therefore, we predict that key environmental drivers of FCL are those that change operationalspecies richness from low to intermediate levels.

In SI 3, we repeated our analysis with empirical food webs with higher resolution. Although the amount of such food web data was small (11 food webs), operational species richness positively correlated with FCL, and the saturating functional form of operational species richness predicted empirical FCL.

### 3.2 Operational species richness modulates environmental effects on FCL

Our simulation shed light on how environmental factors – resource availability, disturbance, and ecosystem size – dictate FCL through changes in species richness. In the main text, we focused on the effects of disturbance and ecosystem size because our preliminary results suggested very weak correlations between resource availability and FCL (see SI 7 for the analysis including resource availability). Consistent with the previous theory, small and frequently-disturbed ecosystems were unable to maintain long food chains (Fig. 3A). However, our theoretical prediction is unique because the effect of disturbance was contingent on ecosystem size and vice versa. For example, disturbance strongly regulated FCL in small ecosystems, but such disturbance-induced regulation was weak in large ecosystems (Fig. 3C; second and fourth rows inTable S4; one-sided Wilcoxon signed-rank test *p <* 10^−6^, *T* = 465).Similarly, enlarging ecosystem size resulted in enhancing FCL under strong disturbance but such increase in FCL was smaller under weak disturbance (Fig. 3E; sixth and eighth rows in Table S4; one-sided Wilcoxon signed-rank test *p<* 10^−6^, *T* = 465).

**Figure 3:**
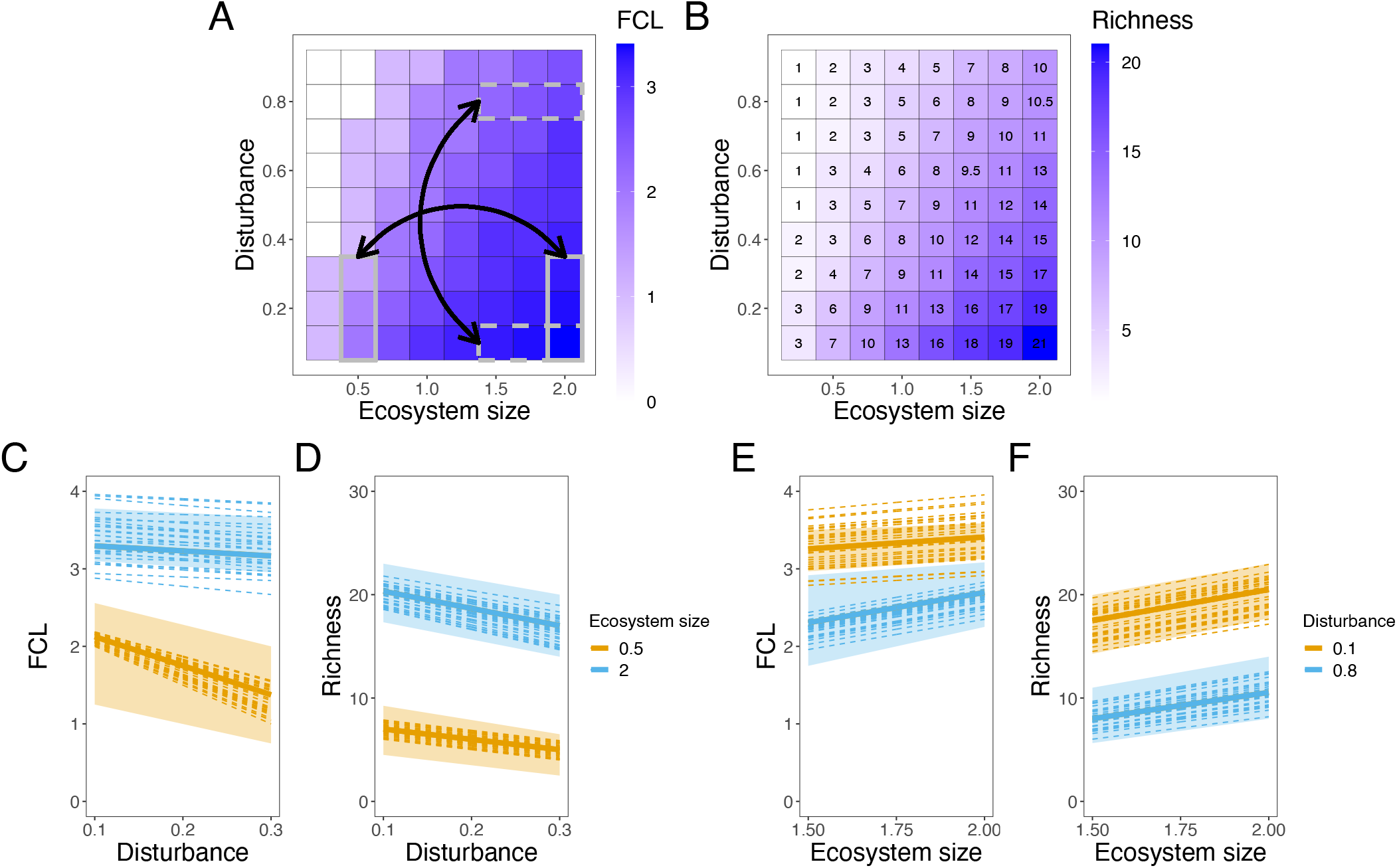
Species richness modulates environmental effects on FCL. A-B: Median values of FCL mean (A) and operational species richness (B) in our simulations over two environmental factors (disturbance and ecosystem size) are shown. On the one hand, operational species richness consistently decreased over disturbance but increased over ecosystem size; see also panels D and F. On the other hand, the two environmental drivers little affected FCL at high operational species richness (i.e., weak disturbance and large ecosystem size). The boxes in panel A represent FCL in panels C (the solid-line boxes) and E (the dashed-line boxes). C-F: Distributions of FCL (C and E) and operational species richness (D and F) are shown. The solid lines represent median values of the fixed effects, the dashed lines include the random effects of different 32-species food web structures, and the shaded areas show 25 % - 75% quantile of the fixed effects. The environmental effects on FCL are clear when operational species richness was low (*<* 15); at high species richness, however, FCL was robust against the changes in the environments. C: Disturbance decreased FCL more in a smaller ecosystem (orange) than in a larger one (blue). D: Disturbance decreased operational richness regardless of ecosystem size. E: Ecosystem size increased FCL more under strong disturbances (blue) than weak ones(orange). F: Operational richness increased over ecosystem size regardless of the frequency of disturbances. See Tables S4 and S5 to compare the slopes in the linear quantile mixed models.

The non-linear FCL-richness association underlay the emergent dual regulation of FCL. As expected, operational species richness decreased over disturbance while increased over ecosystem size (Fig. 3B). In contrast to the pattern of FCL, disturbance regulated operational species richness regardless of the ecosystem size, and vice versa (Figs. 3 D and F). The decoupling of FCL and richness patterns over environmental gradients can be explained by the saturating form of the FCL-richness relationship. For example, when operational species richness exceeded 15, changes in species richness did not translate into FCL changes due to the saturating functional form. Thus, operational species richness modulated how disturbance and ecosystem size affected FCL. These patterns were consistent regardless of the definition of FCL (Fig. S4). Simulations with various structures of *N* -species food webs showed patterns similar to the main text (Figs. S19 – S21). Thefore, operational species richness is a robust modulator between the environmental drivers and FCL.

## 4 Discussion

Our mechanistic exploration highlighted the innate complexity of FCL responses to environmental drivers, offering the potential explanation for the mixed results in previous studies (Sabo et al., 2009, 2010; Takimoto and Post, 2013; Warfe et al., 2013; Young et al., 2013). In particular, the effects of ecosystem size and disturbance were contingent on each other. While previous studies (Sabo et al., 2010; Pomeranz et al., in press) suggest the context-dependent effects through the relationships among potential environmental drivers, our theory does not require such relationships. Instead, the context dependency in our model arose from the nonlinear relationship between FCL and operational species richness (Figs. 2D, and 3A and B). This nonlinear pattern is confirmed by empirical food web data (Fig. 2A), and consistent with previous studies (Martinez and Lawton, 1995; Vander Zanden et al., 1999). Hence, our results suggest the prevalence of “apparent” interactions between multiple environmental drivers, implying that the observed contingency may be a natural outcome of complex food webs. The gradient of operational species richness across environments would be key to resolving inconsistencies in previous FCL studies.

The lack of resource effects (see SI 7 for including resource availability in the main text model, and Fig. S9B for an alternative model) seems counterintuitive because some previous studies report the importance of resource availability (Post and Takimoto, 2007; Doi et al., 2009; Kondoh and Ninomiya, 2009; Takimoto and Post, 2013; Ward and McCann, 2017; Terui and Nishijima, 2019). This mismatch may be attributable to the difference in the number of operational species considered. Some of the above studies analyzed simple systems composed of three or four species (Post and Takimoto, 2007; Doi et al., 2009; Ward and McCann, 2017; Terui and Nishijima, 2019) unlike our theoretical model (32 species). These studies suggest that FCLs are lengthened with increasing resource availability at lower ranges (Post and Takimoto, 2007; Terui and Nishijima, 2019; Ward and McCann, 2017), although excessive productivity may shorten FCL if intraguild predation exists (Post and Takimoto, 2007; Ward and McCann, 2017). It is possible that this prediction is only applicable to simplified food webs with a few species. Another, but not mutually exclusive explanation is that the effect of resource availability could be simply weaker than other environmental drivers. Although Kondoh and Ninomiya (2009) analyzed the effect of resource availability in *N* -species communities, their model does not include ecological processes driven by ecosystem size and disturbance. Our model complemented these components (see also SI 4 for the abundance-based model), finding the lack of noticeable effects of resource availability. These differences may account the contrasting result of resource availability between ours and others.

Like any theoretical research, our results must be viewed with some caution. First, our results might be a unique attribute of the particular food web model we employed. However, we are confident that this is not the case. The saturating effect on FCL appeared regardless of the assembly rules we considered, including random graphs and cascade models (Cohen and Newman, 1985) (see SI 6). Thus, this saturating response of FCL to operational species richness may apply to a broad range of networks. While we did not consider food web models that produce cyclic food webs (e.g., the niche model (Williams and Martinez, 2000)), it is difficult to envision that cycles qualitatively alter our results because the saturating effect appeared even in random graphs. In support of this view, the saturating function accurately predicted FCL in more recent food web data with the higher taxonomic resolution and cyclic food webs (SI 3), which addressed the potential limitation of Cohen’s larger database, i.e., inaccurate estimates of trophic positions due to the low taxonomic resolution. This fact suggests the robustness and generality of our findings since the non-linear relationship was responsible for the contingent effects of environmental drivers on FCL. Second, we did not account for variations in species’ traits other than their trophic positions in a food web. In nature, species may differ in their colonization rates, sensitivities to disturbances, or competitive abilities. Nevertheless, we could reproduce empirical patterns of how FCL changes over the gradients of food web structures. Hence, introducing more complexity to our model will unlikely overturn our conclusions.

Our findings may also provide important insights into the stability-diversity debate (McCann, 2000). “May’s paradox” of community stability has sparked a discussion of how complex webs of interacting species are maintained despite their inherent instability (Gardner and Ashby, 1970; May, 1972; Allesina and Tang, 2012), and previous studies suggest that the non-random species interaction is the key factor promoting stable coexistence (Thébault and Fontaine, 2010; Mougi and Kondoh, 2012; García-Callejas et al., 2023). The present study suggests another non-randomness related to community stability: expanding the “vertical” dimension of biodiversity (Wang and Brose, 2018), or increasing FCL, is more likely to lead to community collapse than increasing “horizontal” biodiversity or adding species to the already-occupied trophic positions. The chain motif is distinctive because species within the chain motif occupy broader trophic positions than others (see Fig. S22 and Table S15). Increasing chain motifs means that species occupy open and higher trophic positions, resulting in longer FCL. However, Figs. 2E and F indicate that chain motifs were unlikely to be prevalent in species-rich communities; instead, colonized species likely occupied trophic positions similar to resident species, forming competitive motifs with the saturating increase in FCL (Figs. 2 A-D). It is reasonable to observe the saturating effect of operational species richness on FCL (Fig. 2). This finding may open an opportunity to link the ongoing diversity-stability debate to FCL research.

Our findings do not mean to downplay the role of stable isotopes in FCL research; instead, these suggest the importance of combining multiple methods to gain deeper insights. Although stable isotopes are applicable to a variety of natural systems, they treat internal food web structure as a “black box.” Emerging molecular techniques, such as (meta-)barcoding of environmental DNA, can investigate the food web structure (Taberlet et al., 2018; Pringle and Hutchinson, 2020). Recent studies use such techniques to assess species’ diet (Deagle et al., 2009), to reconstruct food webs structure (DAlessandro and Mariani, 2021), or to estimate interspecific interaction strength (Ushio et al., 2023). These analyses have the potential to unveil detailed internal food web structures, including species richness and fractions of food web motifs.

In conclusion, this manuscript sheds light on the role of internal food web structure in producing the context-dependency of FCL controls. Uncovering food web structure in nature may prove challenging. However, recent technological advancements have created opportunities to address this issue. In this regard, our theoretical framework serves as a conceptual foundation for future studies that utilize emerging methodologies to investigate food webs: how operational species richness and, to a lesser extent, food web motifs modulate the associations between FCL and environmental drivers. By incorporating the intrinsic complexity of food webs into FCL research, we can potentially resolve contradictory results observed in various ecosystems.

## Supporting information

Supplementary analysis

