## Supplementary analysis for "Food web complexity modulates environmental impacts on food chain length"

Supporting information

### SI 1 Details of measuring FCL and motifs

#### SI 1.1 Measuring trophic positions and FCL

In the main text, we introduced three definitions of trophic positions and corresponding FCL (short, mean, and long). In this subsection, we explain the details of how we measured the trophic positions and FCL.

FCL mean accounts for quantitative information on prey-predator interactions. Mathematically, trophic positions in this case are given by the solution of the following simultaneous linear equations:

$$\tau_i = 1 + \sum_{j=1}^{N} A_{ji}\tau_j, \quad i = 1, \ldots, N, \tag{S1}$$

where $\tau_i$ is species $i$'s trophic position and $A_{ij}$ represents a *weighted* adjacency matrix of a focal food web:

$$\begin{cases} A_{ji} > 0 & \text{if } i \text{ eats } j \\ A_{ji} = 0 & \text{otherwise} \end{cases} \tag{S2}$$

and

$$\sum_{j=1}^{N} A_{ji} = \begin{cases} 0 & \text{if } i \text{ is basal species} \\ 1 & \text{otherwise.} \end{cases} \tag{S3}$$

The above simultaneous linear equations always have unique solutions when a food web is acyclic. In cyclic food webs, the unique solution may not exist. In such cases, we dealt FCL mean as “not available”. The unique solution exists in the simultaneous linear equations if and only if the rank of $N \times N$ coefficient matrix of the linear equations is $N$. We calculated the rank using rankMatrix() function in Matrix library version 1.5-3 (Bates et al., 2022). When the simultaneous linear equations have a unique solution, it was obtained by solve() function in R.

In FCL short and long (see SI 2), we used binary information of prey-predator interactions (whether interactions exist or not) in a directed network. If trophic positions are determined by the shortest-path length from the basal species, we measured the shortest distance from the basal species to a focal species in an adjacency matrix of a food web using the distance function in igraph library (Csardi and Nepusz, 2006) in R. If the trophic levels are determined by the longest-path length, on the other hand, we measured the shortest distance from the basal species to a focal species in an adjacency matrix whose elements are negative values of an original food web. This method worked because we measured FCL long in acyclic food webs.

##### SI 1.2 Meauring motifs fractions

The triad_census() in igraph provides numbers of 16 motifs in networks. We removed 3 of them that include isolated nodes following previous studies (Stouffer et al., 2007; Borrelli, 2015; Monteiro and Faria, 2016). Because species richness and connectance affect the number of motifs, we used the fractions of motifs (the number of the focal motif over the sum of 13 motifs).

#### SI 2 Alternative definitions of FCL

In this section, we perform our analysis in the main text while changing the definition of FCL. Depending on how to measure the trophic positions, we have two alternative definitions of FCL: FCL long and short, respectively. While FCL mean in the main text reflects the quantitative difference in predations, these two alternative FCL use binary prey-predator interactions (i.e., whether trophic interactions exist or not). In all definitions of FCL, operational species richness and the fraction of the three-species chain motifs positively correlated with FCL (in empirical food webs: Fig. S1, and in simulations: Fig. S2). In addition, the two subsections below show that operational species richness still modulated the environmental effects on FCL when we used FCL short and long.

##### SI 2.1 FCL long

One alternative way to define trophic positions is to define one plus the longest-path length from basal species. When FCL is given by the trophic positions defined above, we call it FCL long. Although we cannot measure this FCL in cyclic food webs, FCL long is always given by the longest-path length from a basal species to a top predator (species without their predators).

As in the main text, Figs. S3A-D shows that the functional responses of operational species richness and the fraction of the three-species chain motifs to FCL were the saturating functions and quadratic ones, respectively (see also Table S3). In addition, the coefficients of the best models suggested that species richness had a dominant effect on FCL. The fraction of the chain motifs was lower than expected in the random graph ( Figs. S3E and F). Although weaker disturbances and larger ecosystems increased operational species richness (Fig. S4C), this did not reflect in the enhancement of FCL long when species richness exceeded 15 (Fig. S4A). Therefore, operational species richness also modulated the environmental effects on FCL long.

##### SI 2.2 FCL short

The other alternative definition of trophic positions is quantified by one plus the shortest-path length from a casal species. We call FCL defined by such trophic positions FCL short. This FCL can be measured both in cyclic and acyclic food webs.

Figs. S5A and B show that operational species richness again had the saturating effect on FCL both in the empirical data and our simulation. The fraction of the chain motifs, however, had a linear effect on FCL

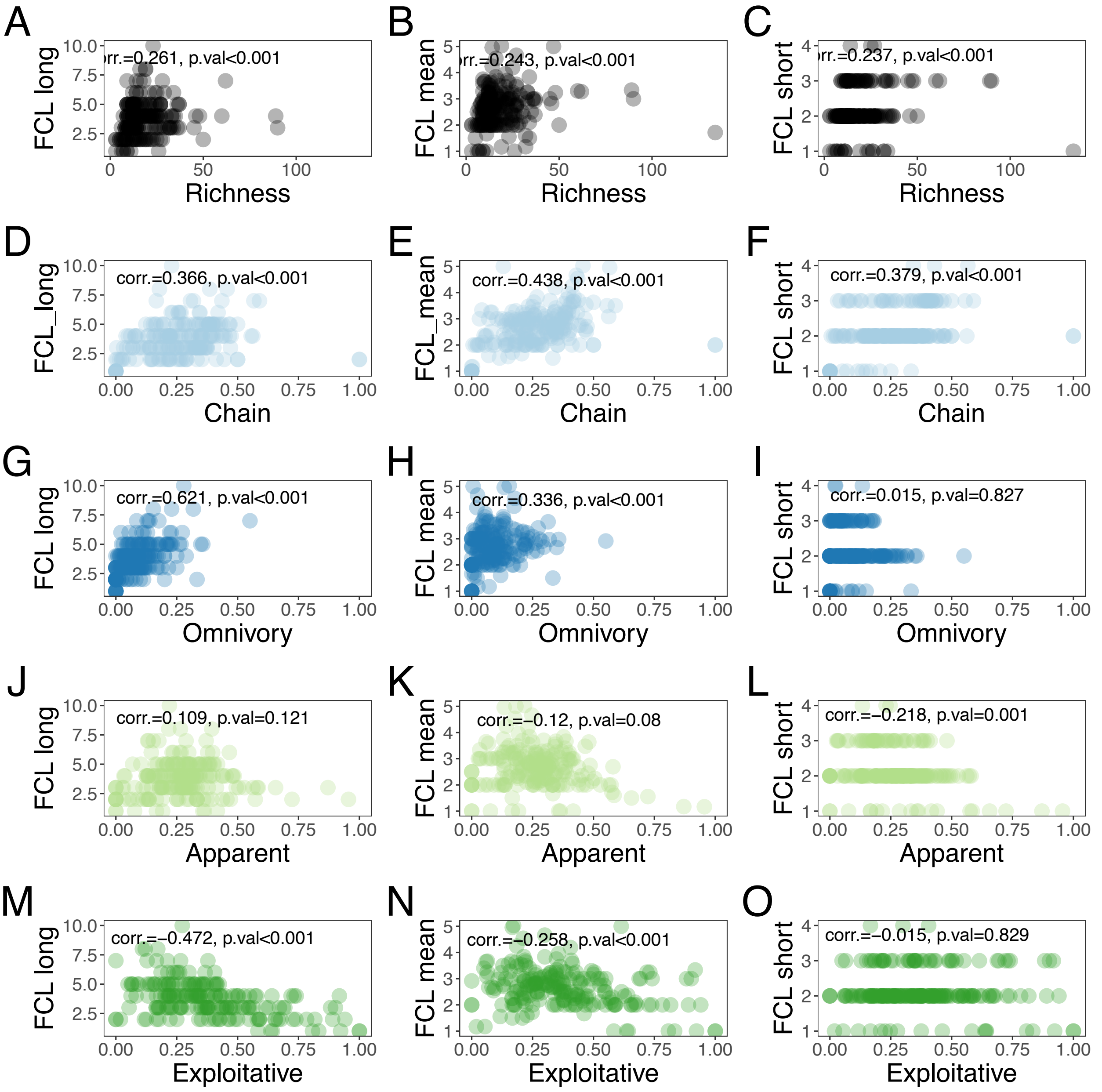


Figure S1: Correlations in empirical food webs

Spearman correlation coefficients between FCL and species richness (A-C) or the fractions of food web motifs (chain: D-F, omnivory: G-I, apparent competition: J-L, and exploitative competition: M-O) in the empirical food webs are shown. The correlation results were obtained from cor() function in R and we removed data of operational species richness $< 3$ because we cannot define the motifs there. We measured three types of FCL depending on how we define trophic positions. On the left column (FCL long), we define trophic positions as one plus the longest-path length from the basal species. On the center column (FCL mean), species' topic levels are given by one plus mean of prey species' trophic positions. On the right column (FCL short), trophic positions are given by one plus the shortest-path length from the basal species. The p-value of the correlation test is also shown on each panel.

in the empirical food webs and a quadratic effect on FCL in our simulations, respectively (Figs. S5 C and D, respectively). Nevertheless, the coefficients in the best model suggested that operational species richness was the main contributor to FCL (Table S3) because the chain motif could increase FCL at most 2.12 while operational species richness did 2.85. Although weaker disturbances and larger ecosystems increased FCL, these environmental effects were masked when operational species richness exceeded 16. The saturating effect of operational species richness on FCL produced the inter-dependent effects of disturbance and ecosystem size(Figs. S4B and D). Taken together, our results in the main text were maintained when we used FCL short or long instead of FCL mean.

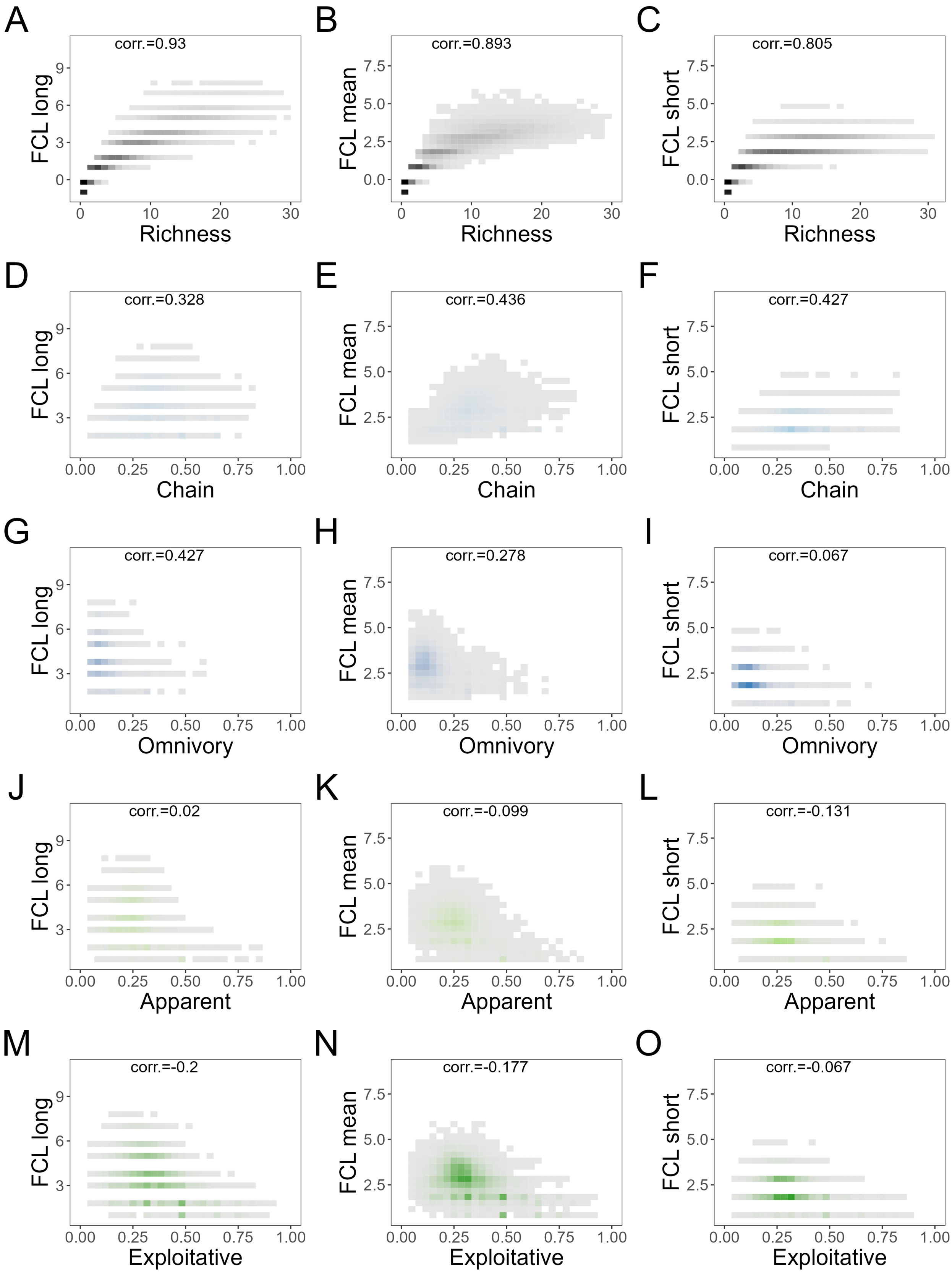


Figure S2: Correlations in food web simulations

Correlation plots similar to Fig. S1 are shown. Because of the large number of simulation data, p-values are always small and omitted. The darker, bluer, or greener areas represent more data than gray areas.

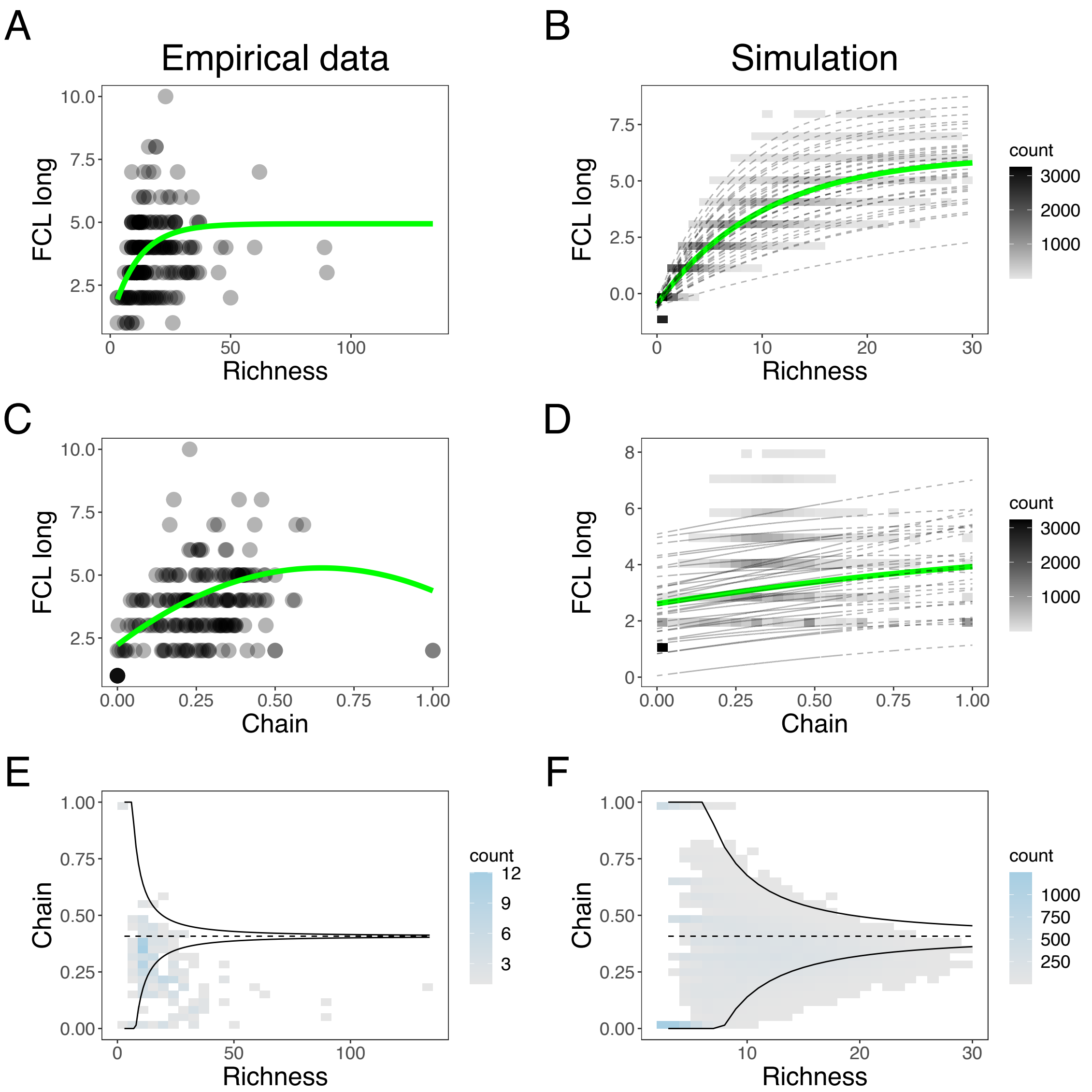


Figure S3: Species richness and the chain motif relate to FCL long

Similar to Fig. 2, but the figures here represent the result of FCL long. Note that panel E on this figure is identical to Fig. 2E because the definition of FCL does not affect either operational species richness or the fraction of the chain motifs in the empirical data.

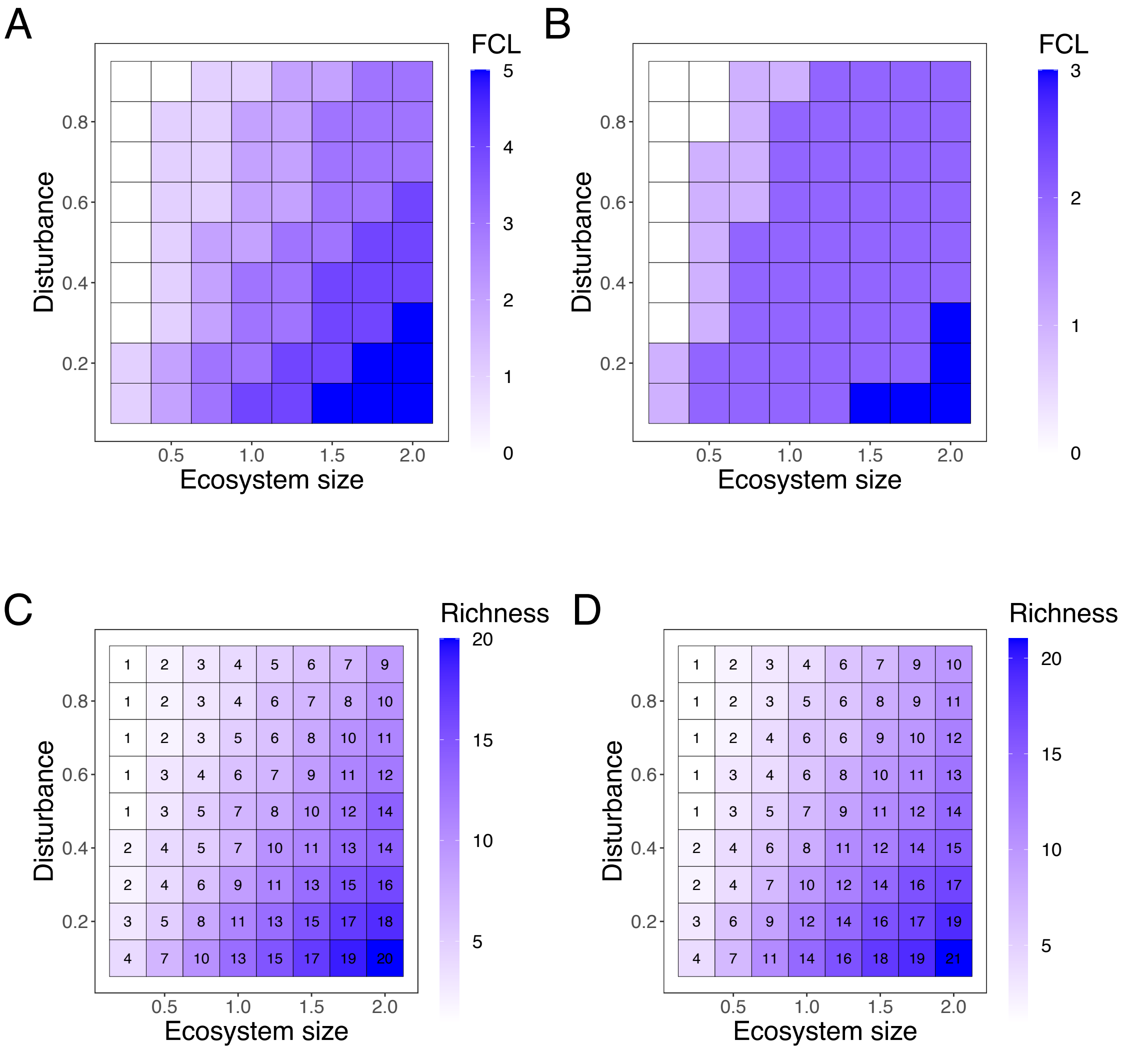


Figure S4: Operational species richness modulates environmental effect on FCL long and short

A and B: Similar to Fig. 3A, median values of FCL in our simulation over disturbance and ecosystem size are shown. However, FCL is defined by FCL long (A) or FCL short (B), respectively, in this figure. C and D: Similar to Fig. 3B, median values of operational species richness are shown over the two environmental values when FCL is defined by FCL long (C) and FCL short (D).

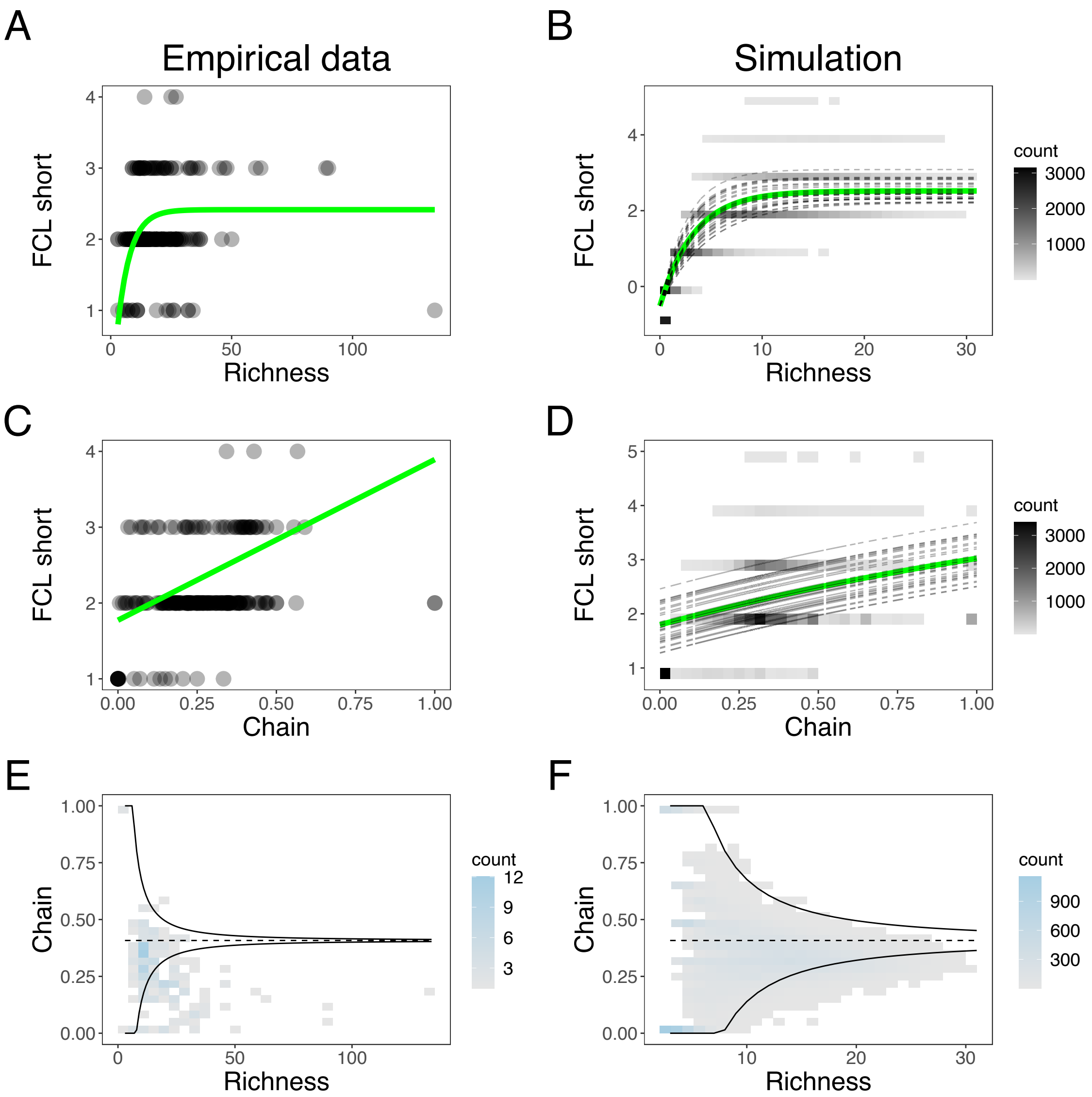


Figure S5: Operational species richness and the chain motif relate to FCL short

Similar to Fig. 2, but the figures here represent the result of FCL short. Note that panel E on this figure is identical to Fig. 2E because the definition of FCL does not affect either species richness or the fraction of the chain motifs in the empirical data.

#### SI 3 Another empirical database

Empirical food webs in Cohen (2010) have low resolutions in two senses: the low resolution of nodes (e.g., a node in WEB1 in the database represents carnivore fish), and low resolution of edges (i.e., many trophic interactions are binary). This could be a potential problem in the analyses in the main text because FCL and trophic positions can change. In this section, we analyzed other 11 empirical food webs containing higher taxonomic resolution and quantitative edges (Angelini and Agostinho, 2005; Angelini et al., 2006, 2010; Angelini and Vaz-Velho, 2011; Angelini et al., 2013; Bascompte et al., 2005; Christian and Luczkovich, 1999; Cruz-Escalona et al., 2007; Torres et al., 2013) obtained from Web of Life (`https://www.web-of-life.es/map.php?type=7`). As of April 18th, 2023, this food web database contains 33 food webs. We removed food webs with binary edges and those focusing on bipartite networks (i.e., FCL is 1 and the chain and omnivory motifs do not exist) from the analysis, resulting in 11 remaining food webs. Because FCL short and long do not rely on quantitative edges, we only analyzed FCL mean in this section.

Fig. S6 shows how species richness or the fractions of food web motifs correlate with FCL in these new food webs. FCL mean positively correlated with species richness, consistent with the results in the main text. Within the four food web motifs, only the chain motif positively correlated with FCL, but the correlations of all four motifs were not statistically significant, probably due to the small amount of food web data.

We also determined whether species richness and the fraction of the chain motif predicted FCL mean in the new database. The regression analysis and model selection with the new empirical data, however, would not be meaningful due to the small amount of data. Instead, we compared the actual FCL mean in the new database and the predicted ones by the best model in Cohen's database (the second row in Table S3, Fig. S7), where operational species richness and the fraction of chain motifs have a saturating and a quadratic effect on FCL, respectively. The model predictions were close to the actual FCL mean, except for two data (FCL is approximately 3.8 and 4.5, respectively). The failure to fit to these data may reflect the small number of long FCL mean data in Cohen's database (see Fig. 2A); i.e., the model cannot be trained to fit long FCL. Taken together, the results of this section do not contradict the main text where empirical food webs have lower resolution.

#### SI 4 Implementing population dynamics

While our model in the main text implements the dynamics of the presence and absence of each species, some previous studies implemented population abundance. This section shows another version of the model in which population dynamics are implemented. As we shall see, however, implementing the population dynamics shows similar results to the main text.

This alternative model uses ten adjacency matrices of food webs used in the main text. However, we introduced several parameters to implement the population dynamics: for each species $i$, we need growth rate $r_i$, self-regulation intensity $s_i$, consumption rate on prey $j$ $a_{ji}$, and conversion rate $c_i$. The population dynamics

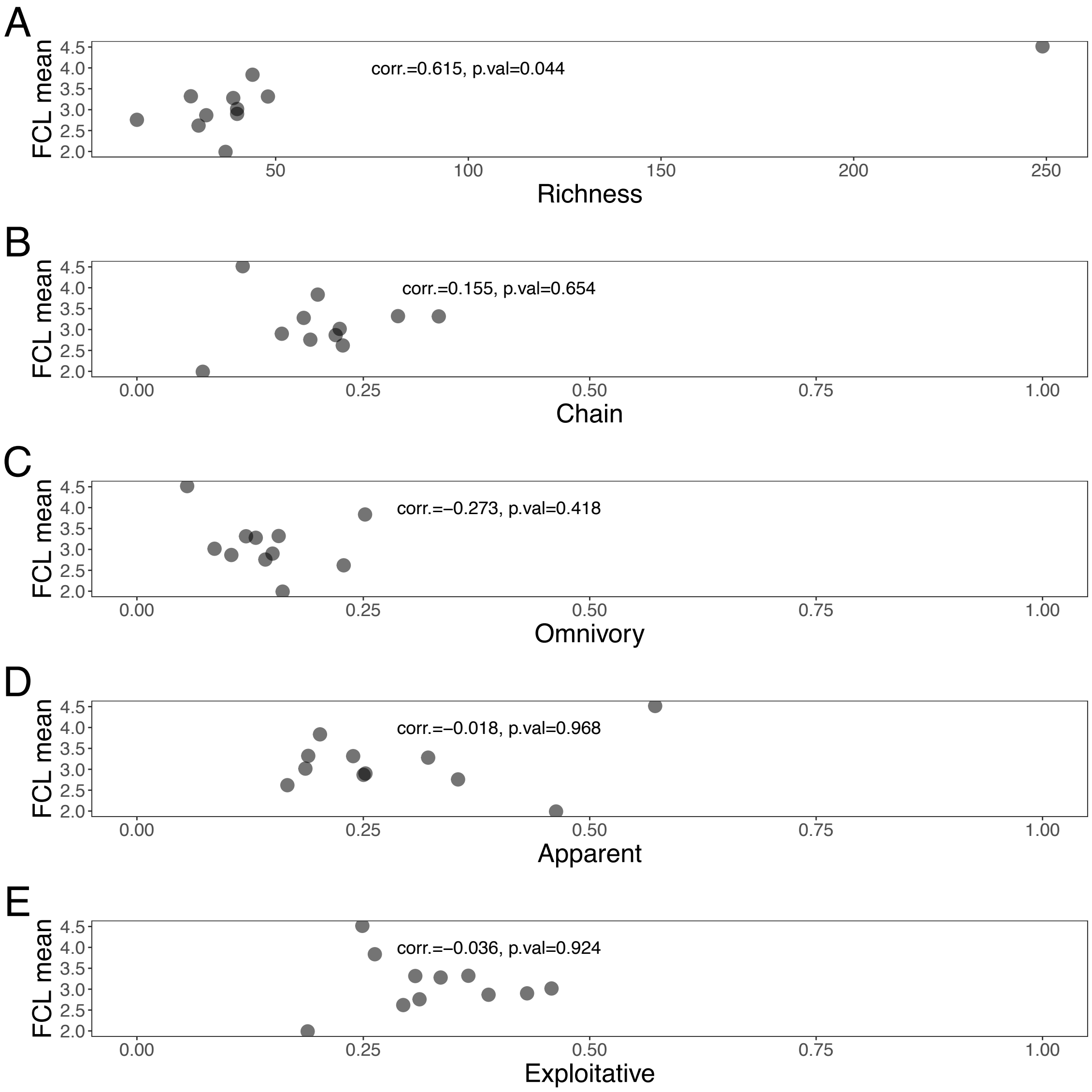


Figure S6: Correlations with FCL in the new empirical food webs

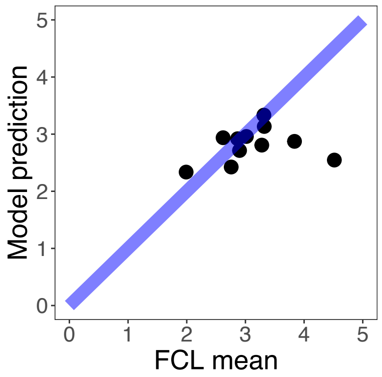


Figure S7: Predicting of FCL mean of the new data

FCL mean in the new empirical food web data are compared with the model prediction from empirical food web data in Cohen (2010) (Empirical FCL mean in Table S3). The blue line indicates the cases where the model can perfectly predict FCLs. R-squared of Model Prediction ∼ FCL mean was 0.96 in lm() function of R.

of species $i$ then is written as follows:

$$\frac{dX_i}{dt} = X_i(r_i - s_i x_i + c_i \sum_{j \neq i} a_{ji} X_j - \sum_{k \neq i} a_{ik} X_k) \tag{S4}$$

where $X_i$ represents abundance of species $i$.

The parameter values were sampled or determined following the models in previous studies (Kondoh and Ninomiya, 2009; Mougi, 2022). The growth rates $r_i$ differed in basal and non-basal species; basal species has a positive growth rate depending on resource availability $R$ while a non-basal species always has a negative growth rate so that they cannot survive without prey (Kondoh and Ninomiya, 2009):

$$\begin{aligned} r_i &\sim \mathcal{N}\left(R, 0.5^2\right) \quad && i = 1, \ldots, B \\ r_i &= -0.01 && i = B+1, \ldots, N \end{aligned} \tag{S5}$$

where $\mathcal{N}(\mu, \sigma^2)$ represents a normal distribution whose mean and standard deviation are $\mu$ and $\sigma$, respectively. Resource availability $R$ ranged from 1 to 125. The consumpition rates $a_{ij}$ range from 0.1 to 1.0 when species $i$ consumes $j$; otherwise, $a_{ij}$ was set zero as in Kondoh and Ninomiya (2009). The conversion rate was fixed across species, $c_i = 0.2$, as in Mougi (2022). In previous studies (Kondoh and Ninomiya, 2009; Mougi, 2022), self-regulation intensity was set to either 0 or 1.0. In our case, we set $s_i = 0.1$ so that population sizes are bounded, but growth rates are larger than the case $s_i = 1.0$. This is because our model included disturbance that randomly removes species, while the two previous studies do not.

The effects of disturbance and ecosystem size were implemented as in the main text: the extinction and colonization rates of species, respectively. When a disturbance occurs, a randomly chosen species in the focal patch goes extinct, while migration increases a randomly chosen species' abundance by 0.1. The dynamics of Eq (S4) were implemented by deSolve package version 1.34 (Karline Soetaert et al., 2010) in R using rk4 method. While solving Eq (S4), species are randomly either removed from or introduced to the patch. The waiting time for such stochastic events followed the exponential distribution whose mean was the inverse of the sum of all stochastic events' rates (i.e., $N\times$extinction rate$+$local species richness$\times$colonization rate). The extinction rate and colonization rate ranged between $10^{-4}$ and $10^{-1}$ as proxies for the intensity of disturbance and ecosystem size, respectively. This reflects the stochastic changes in the presence-absence data of species due to disturbance and colonization in the main text.

We measured only FCL mean in this model because the rest two definitions do not incorporate population abundance. FCL mean was measured at the end of each simulation. The weight $A_{ji}$ in Eq (S1) reflects both the consumption rates and the prey abundance at the end of simulations in this case. Mathematically, the weight is given as follows:

$$A_{ji} = \frac{a_{ji} X_j^*}{\sum_{k \neq i}^{N} a_{ki} X_k^*} \tag{S6}$$

where $X_j^*$ represents the population abundance of species $j$ at the end of the simulation

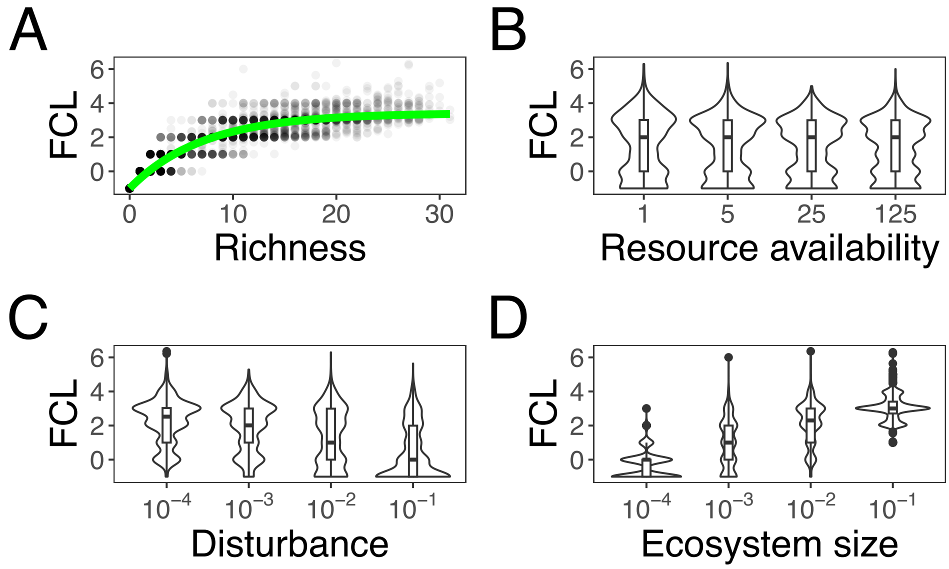


Figure S8: Changes of FCL mean in the model with population dymamics

A: FCL mean shows a saturating pattern over operational species richness. The green curge represents the fit to the simulation data: FCL $= 4.22 \times \{1 - \exp(-0.14 \times \text{richness})\} - 1$. B - D Violin plots of FCL mean over resource availability, disturbance, and ecosystem size.

Fig. S8 shows how FCL changes over operational species richness, resource availability, frequency of disturbance, and ecosystem size in 3200 simulation data. As the results in the main text, the operational species richness has a saturating effect on FCL. While resource availability were not correlated with FCL (Spearman correlation coefficient: $-0.05$, p-value 0.0102), disturbance and ecosystem size were negatively (Spearman correlation coefficient: $-0.43$, p-value $< 10^{-4}$) and positively (Spearman correlation coefficient: 0.76, p-value $< 10^{-4}$) correlated with FCL, respectively. In addition, Fig. S9 shows how FCL and species richness changed over disturbance and ecosystem size. Because of the functional form, disturbance and ecosystem size affected FCL at low operational species richness but did little at high (approximately 15 or more) operational species richness. Therefore, the inclusion of population dynamics did not qualitatively affect the conclusion in the main text (see Fig. 3).

#### SI 5 Alternative implementation of ecosystem size

In the main text, we assumed that species are more likely to colonize in a larger patch (i.e., the target effect). However, Ward and McCann (2017) implemented alternative ecosystem size's effect: decreasing predation rates due to lower encounter rates. In this section, we implemented this idea in our model. Because our model in the main text simulated the dynamics of the presence-absence of species, we assumed that predation pressure was lower in a larger ecosystem and prey species had lower extinction rates due to predation (i.e., lower $c_i$). In

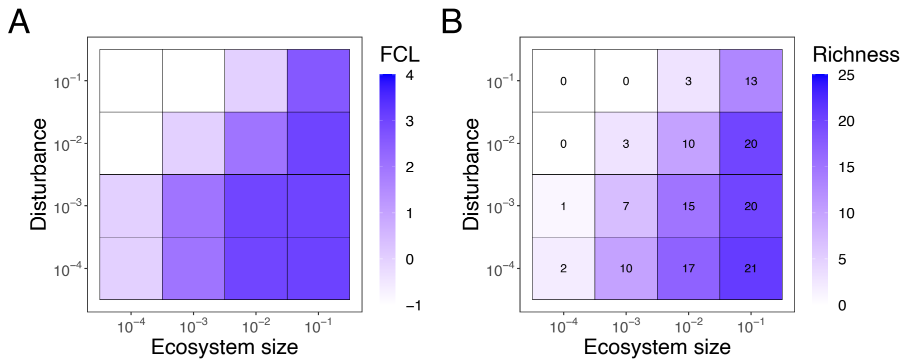


Figure S9: Heatmaps of median FCL mean and operational species richness with population dynamics

Median values of FCL mean (A) and operational species richness at given extinction and colonization rates (proxies for disturbance and ecosystem size, respectively) are shown.

other words,

$$c_i = 1 - \text{ecosystem size} \tag{S7}$$

where ecosystem size ranged from 0.1 to 0.9. Here, we only changed the effect of ecosystem size, and the details of the model (i.e., food web structure, parameters, number of replicates) are as explained in the main text.

Figs. S10–S12 show correlation heatmaps (A), how FCL changes over operational species richness (B) or the fraction of chain motifs (C), and how disturbance and ecosystem size change FCL (D) and operational richness (E). The model selection (Tables S12 – S14) indicates that operational species richness has a saturating effect on FCL, while the fraction of chain motifs has either a quadratic (FCL long and mean) or saturating (FCL short). In addition, due to the saturating effect, operational species richness modulated the relationship between the environmental drivers and FCL. Therefore, the variation in the implementation of ecosystem size did not change the main findings in the main text.

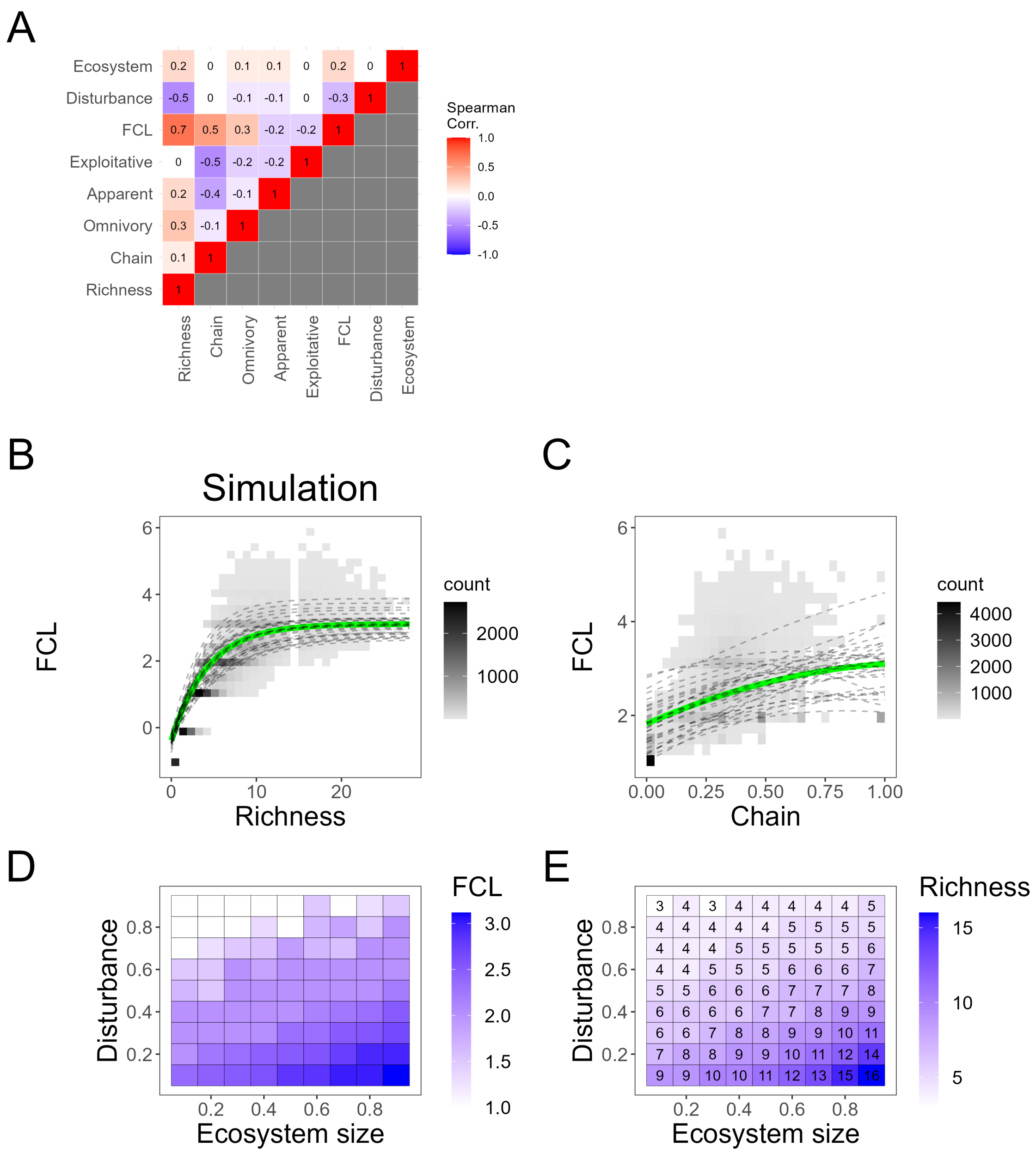


Figure S10: FCL mean in alternative ecosystem size's effect

A: Spearman correlation coefficients are shown. FCL positively correlates with operational species richness and the fraction of chain motifs in realized food webs. B and C: Operational species richness and the fraction of chain motifs are fitted to the saturating and quadratic functions, respectively. The green curves represent $\mathrm{FCL} = 3.50 \times (1 - \exp(-0.23 \times \mathrm{richness})) - 0.93 \times \mathrm{chain}^2 + 2.21 \times \mathrm{chain} - 1$. On panel B, we fixed the fraction of chain motifs as its mean value, while we used mean richness on panel C. D and E: median FCL and operational species richness over two environmental drivers (disturbance and ecosystem size) are shown. In this figure, larger ecosystems have lower extinction rates due to predation, while those in the main text have higher colonization rates.

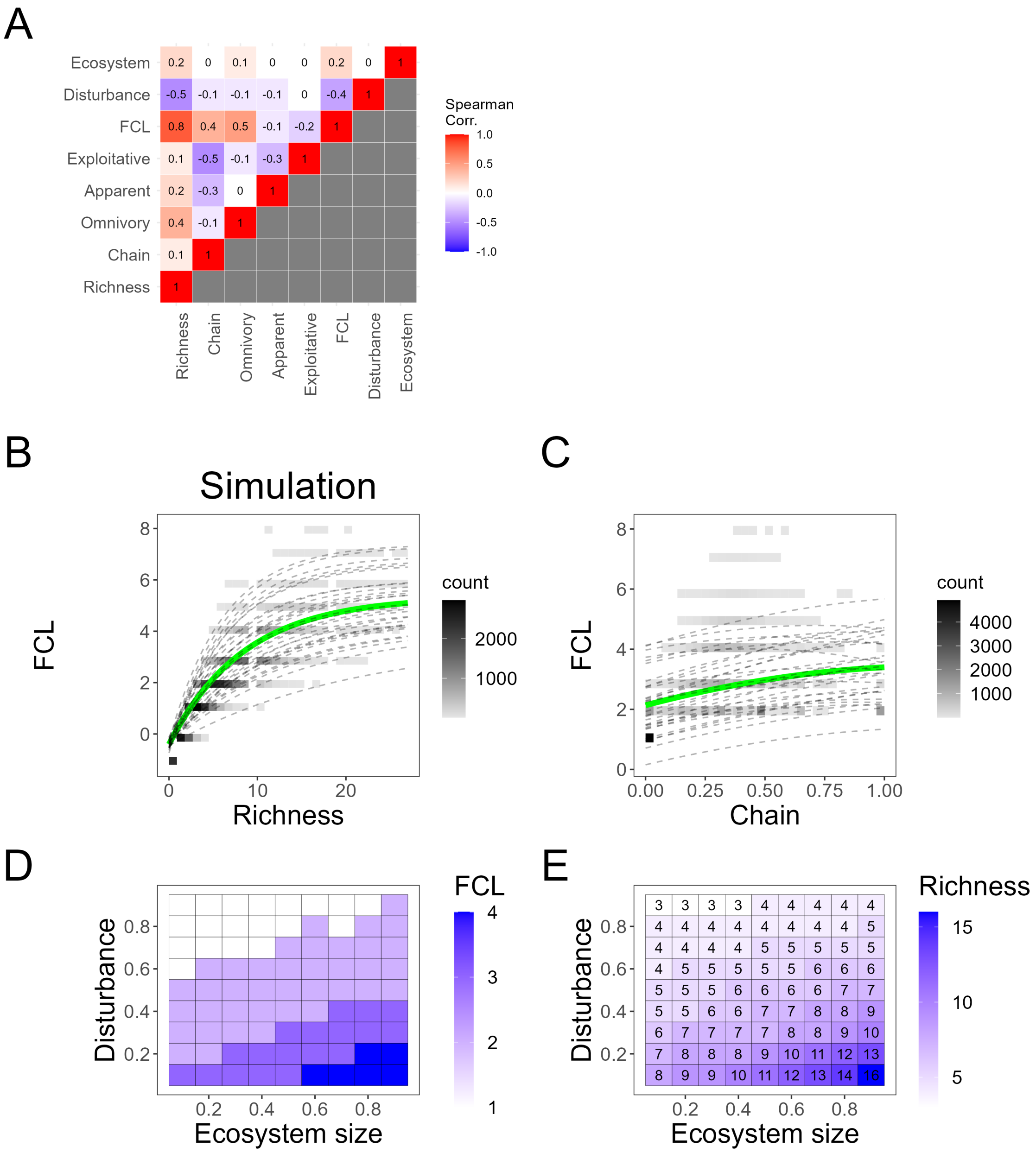


Figure S11: FCL long in alternative ecosystem size's effect

Similar to Fig. S10, but this figure measures FCL long with the alternative implementation of ecosystem size. The green curves on panels B and C represent $\text{FCL} = 5.74 \times (1 - \exp(-0.102 \times \text{richness})) - 0.84 \times \text{chain}^2 + 2.10 \times \text{chain} - 1$.

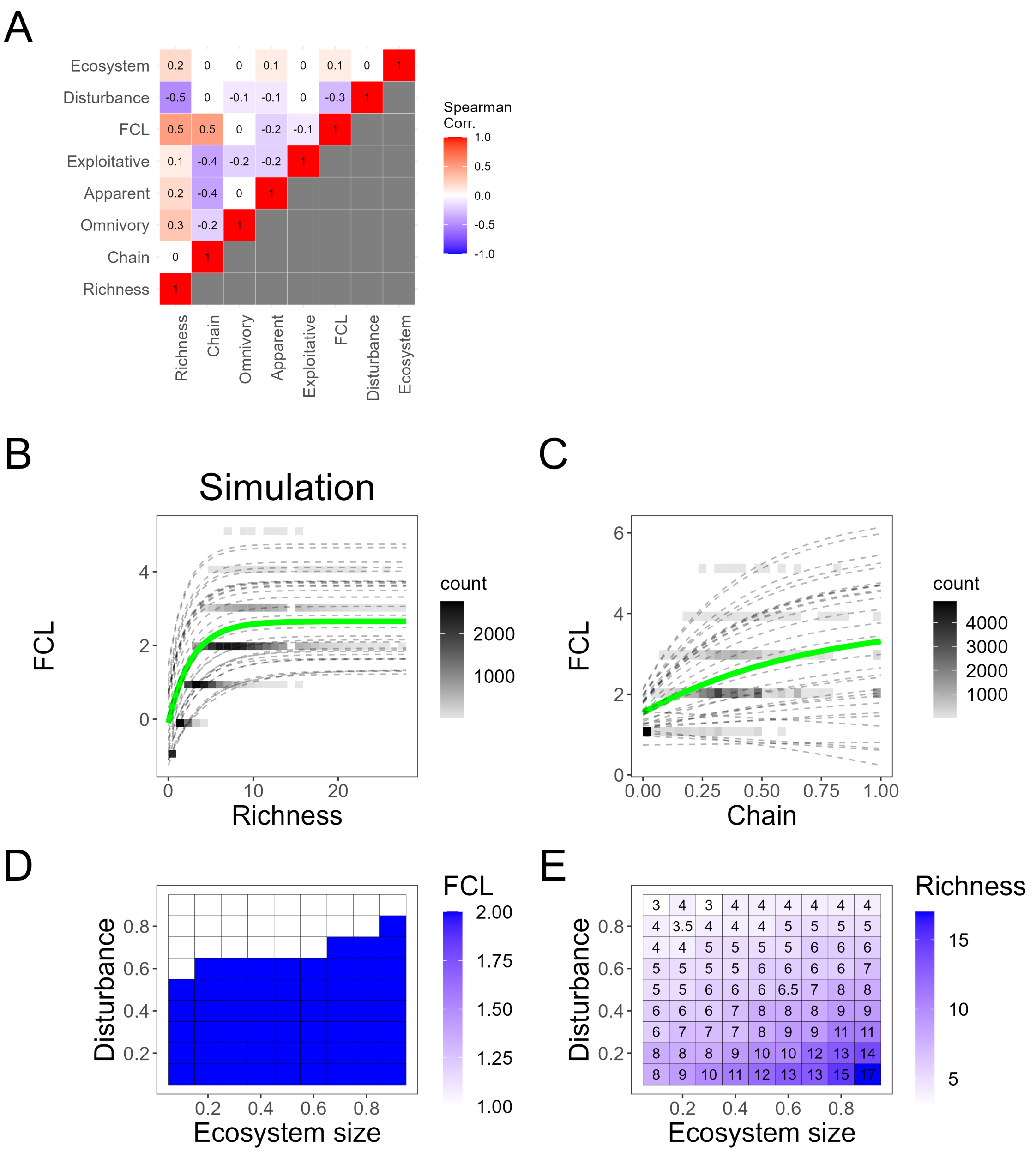


Figure S12: FCL short in alternative ecosystem size's effect

Similar to Fig. S10, but this figure measures FCL short with the alternative implementation of ecosystem size. The green curves on panels B and C represent
$\text{FCL} = 2.75 \times (1 - \exp(-0.36 \times \text{richness})) + 2.34 \times (1 - \exp(-1.40 \times \text{richness})) - 1.$

### SI 6 Two baseline models

In this section, we analyze two types of baseline graphs to deepen our understanding on the relationships among FCL, operational species richness, and the four food web motifs. First, we consider random graphs where an edge between a pair of nodes is assigned with probability $p$. We set $p = 0.11$ so that the expected connectance is identical to food webs produced by the preferential prey model (see Section 2.3). As we set the maximum number of species in our simulation $N = 32$, we also generated random graphs composed of 32 nodes. We generated 30 such random graphs using erdos.renyi.game() function in igraph package (Csardi and Nepusz, 2006) and measured FCL and the fraction of the food web motifs. Additionally, we subsampled $n$ ($n = 3, 4, \ldots 31$) nodes from each full random graph five times and measured FCL and the fractions of motifs to generate variation in operational species richness. In total we obtained $30 \times (29 \times 5 + 1) = 4380$ random graphs.

One benefit of such random graphs is that we can analytically calculate the probability distribution of each motif fraction in a random graph (Fig. S13). Suppose we choose three species from a random graph. This subgraph contains at most six trophic interactions between species, each of which realizes with probability $p$. Then, we can easily calculate the probability that the focal subgraph represents each motif. For example, the three-species subgraph represents the food web motifs with the following probabilities: the chain motif $q_1 = 6p^2(1-p)^4$, the omnivory motif $q_2 = 6p^3(1-p)^3$, the apparent competition motif $q_3 = 3p^2(1-p)^4$, and the exploitative competition motif $q_4 = 3p^2(1-p)^4$. Importantly, three types of subgraphs are removed from motifs; these subgraphs include one or more species that do not interact with others (i.e., isolated species in a subgraph, see SI 1.2). Then, the probability that a randomly chosen three-species motif composes the chain motif is as follows:

$$f_1 = \frac{q_1}{1-Q} \tag{S8}$$

where $Q$ represents the probability that the three-species subgraph includes isolated species:

$$Q = \underbrace{(1-p)^6}_{\text{three isolated species}} + \underbrace{6p(1-p)^5 + 3p^2(1-p)^4}_{\text{one isolated species}}. \tag{S9}$$

In other words, $1-Q$ represents the probability that randomly chosen three species compose a motif. Similar to $f_1$, we can calculate the probability that a randomly chosen motif represents another food web motif as

$$f_i = \frac{q_i}{1-Q}. \tag{S10}$$

Now the number of motifs in a random graph can be modeled by the multinomial distribution. The expected

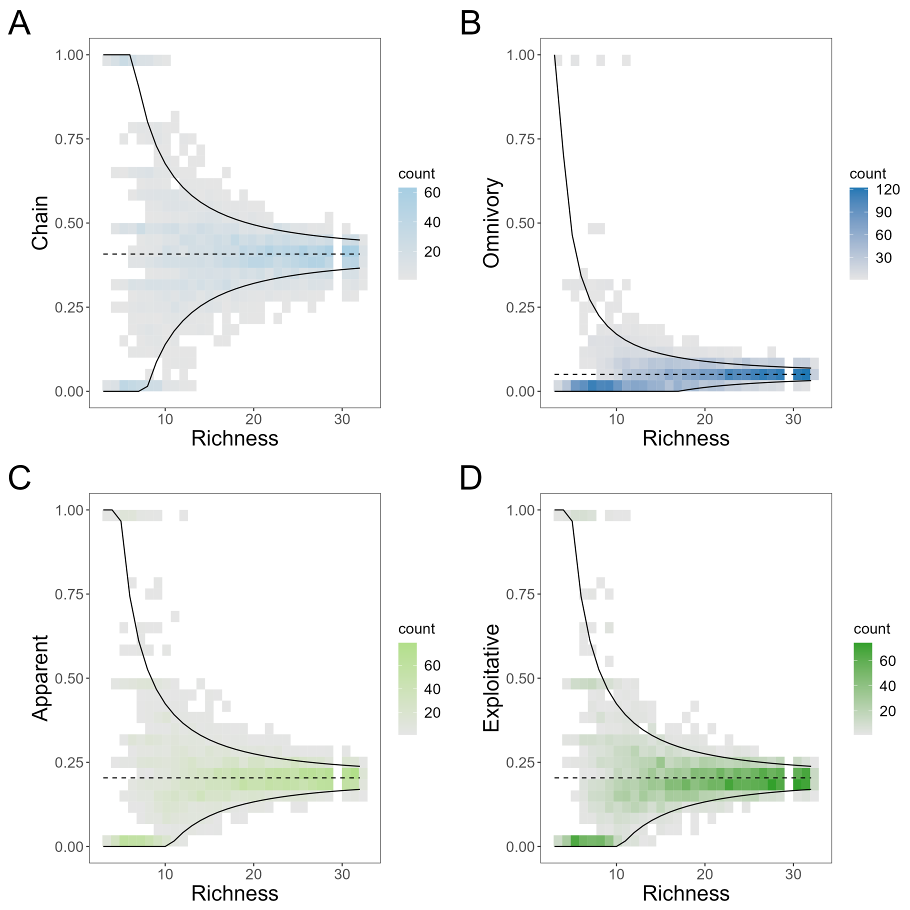


Figure S13: Fractions of motifs in random graphs

: The fractions of the chain motifs (A), the omnivory (B), the apparent competition (C), and the explorative competition (D) over species richness in the random graphs. The dashed lines represent the expected fractions and the solids lines show expected fractions $\pm 2\times$ expected standard deviations.

fraction of motif $i$ and its standard deviation are given as follows:

$$\mu_i = \frac{f_i M}{M} = f_i, \tag{S11a}$$

$$\sigma_i = \frac{\sqrt{M f_i(1-f_i)}}{M} = \sqrt{\frac{f_i(1-f_i)}{M}}, \tag{S11b}$$

where $M$ is the number of motifs in a random graph. Notably, the number of motifs in a random graph is a random variable. When a random graph has $m$ species, the expected number of motif is as follows:

$$\mathbb{E}[M] = \frac{m(m-1)(m-2)(1-Q)}{6}. \tag{S12}$$

Therefore, the expected fraction of motif is constant over operational species richness while its *expected* standard deviation decrease over operational species richness $m$ in the order of $3/2$.

One disadvantage of the random graph as the baseline is that cyclic food webs can be produced, where FCL long cannot be measured and FCL mean can unrealistically inflate (see Fig. S14B). To overcome this problem, we considered a second baseline model: the (classic) cascade mode (Cohen and Newman, 1985), which is one of the oldest food web models. This model generates only acyclic food webs. The upper-right part of the adjacency matrix of a food web is filled with probability $2p(m-1)/m$ so that the expected connectance is $p$. We set $p = 0.11$ and generated 30 32-species food webs using the cascade model. As we did in the random graph, we subsampled $n$ nodes five times from each full graph produced by the cascade model and measured FCL and the fractions of motifs ($n = 3, 4, \ldots 31$). We call this model another baseline because we did not implemented the ecological dynamics as in the main text (see Section 2.3.2)

The two baseline models showed that species richness positively correlated with FCL in all three definitions (Figs. S14 and S15). In contrast, no food web motifs consistently correlate with FCL in these two models. In addition, the model selection suggested that operational species richness had a saturating effect on FCL except for one case (Table S1). Therefore, the saturating effect of operational species richness on FCL would be robust in many graphs.

Table S1: Model selection in the baseline models

| | $\Delta$ AIC in random graph | | | $\Delta$ AIC in cascade model | | |
|---|---|---|---|---|---|---|
| Functional response | FCL long | FCL mean | FCL short | FCL long | FCL mean | FCL short |
| Saturating | 0 | 0 | 0 | 85 | 0 | 0 |
| Linear | 25 | 96 | 112 | 0 | 5109 | 4963 |
| Quadratic | 2.4 | 15 | 19 | 1205 | 1393 | 1530 |

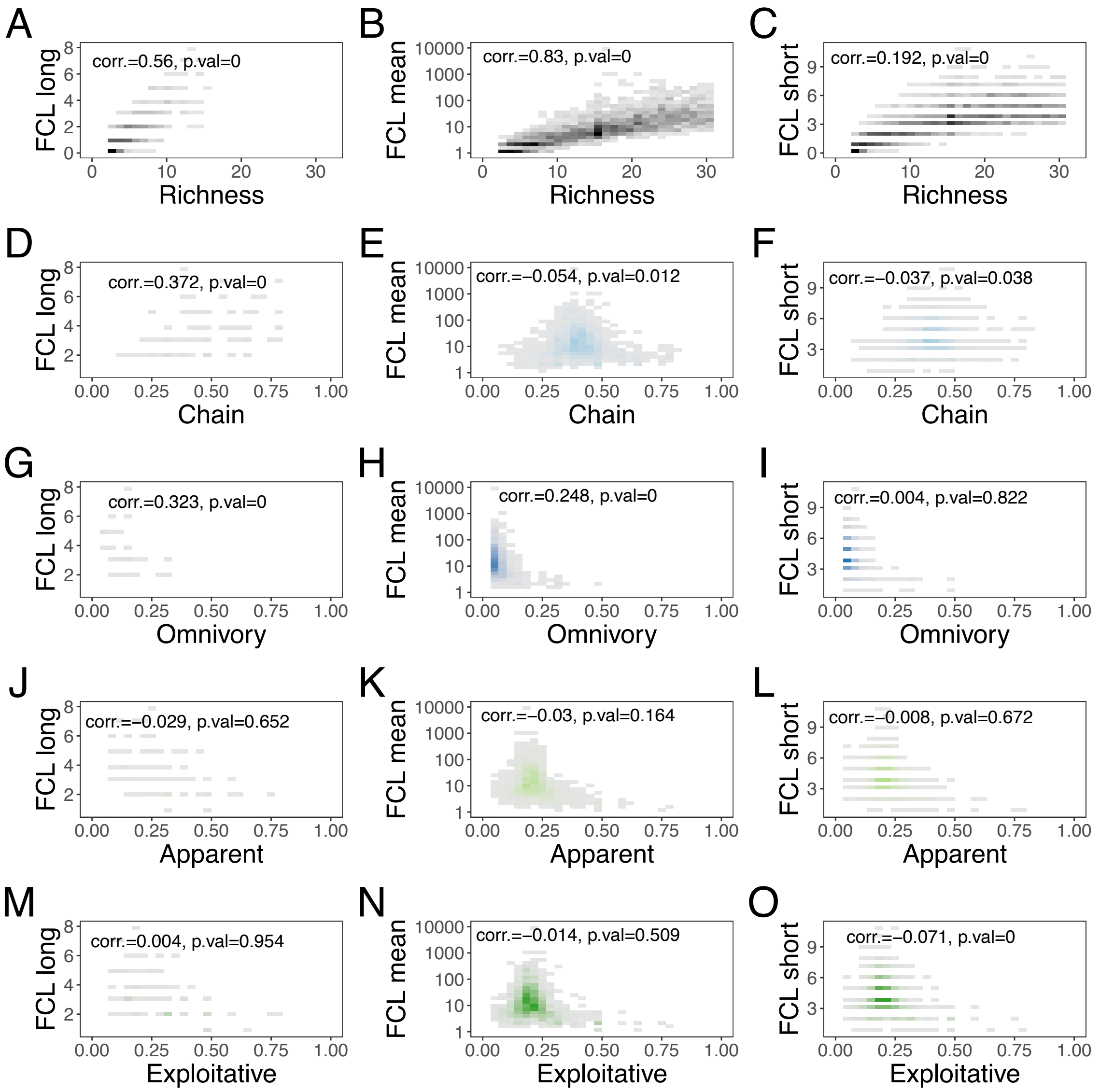


Figure S14: Correlations with FCL in random graphs

Correlations between FCL and species richness or the food web motifs in the random graph are shown. In each panel, Spearman correlation coefficients and p-values are shown on top. The darker, bluer, or greener areas contain more data than gray areas.

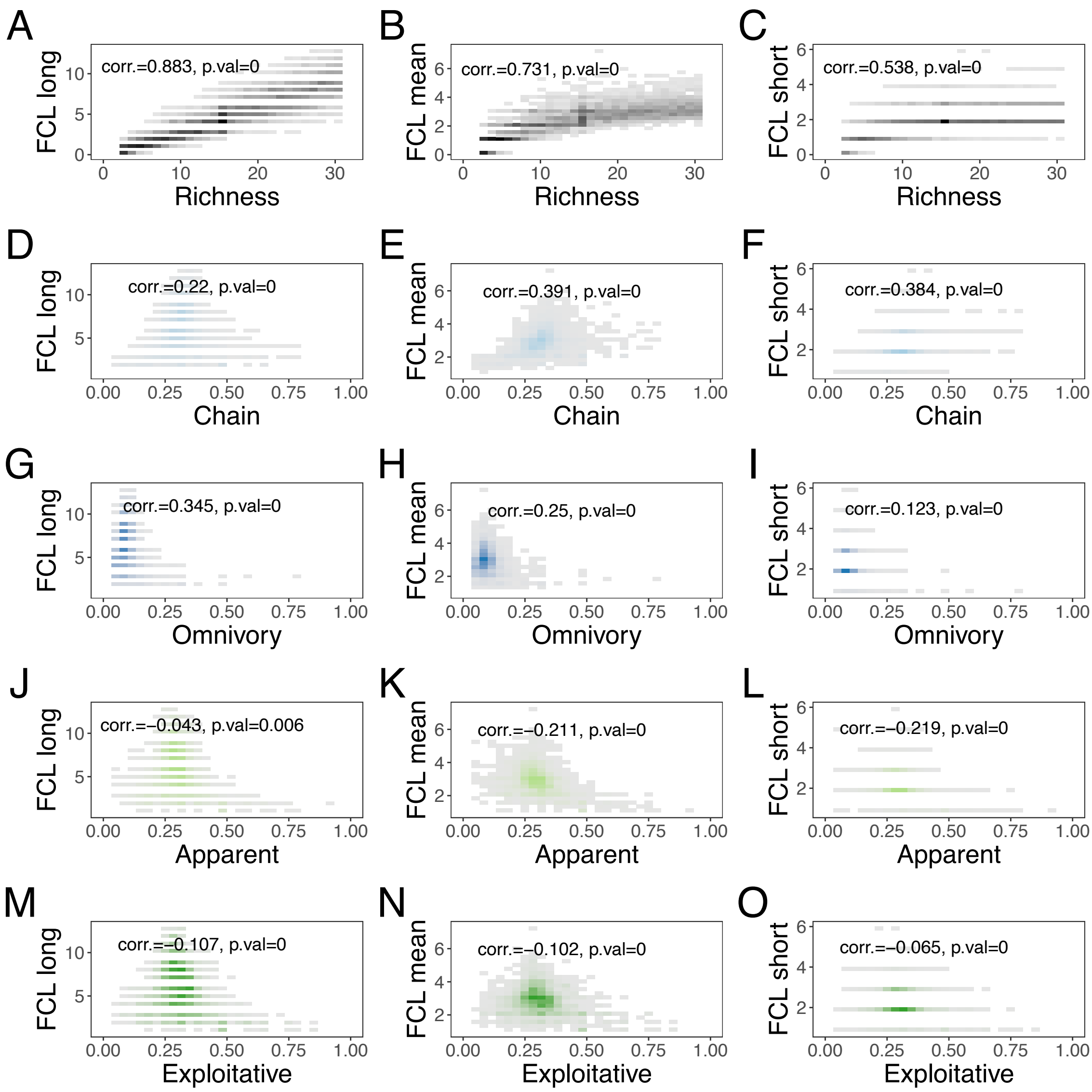


Figure S15: Correlations with FCL in cascade model

Correlations between FCL and species richness or the food web motifs in the cascade model are shown. In each panel, Spearman correlation coefficients and p-values are shown on top. The darker, bluer, or greener areas contain more data than gray areas.

#### SI 7 Anslysis including resource availability

Our preliminary simulations analyzed the effects of resource availability, as well as disturbance and ecosystem size, on FCL and operational pecies richness (see Table S2). In the preliminary analysis, we used the same 32-species food webs in the main text, but we analyzed the sparse parameter values of three environmental drivers while more replicates in each parameter set and food web (100 times per each). The other conditions are identical to the simulations in the main text.

Table S2: Summary of parameters in preliminary simulations

| Symbol | Value | Description |
|---|---|---|
| $N$ | 32 | Number of species |
| $B$ | 4 | Number of basal species |
| $l$ | 113 | Expected number of trophic links |
| $a_i$ | $\in \{0.5, 1, 2\}$ | Maximum colonization rate of basal species |
| $R$ | $\in \{1, 2, 4, 8\}$ | Resource availability |
| $b_i$ | $= a_i$ | Maximum colonization rate of non-basal species |
| $K_i$ | 1 | Number of prey species giving the half-max colonization rate |
| $c_i$ | 0.3 | Maximum extinction rate because of predation |
| $L_i$ | 2 | Number of predator species giving the half-max extinction rate by predation |
| $d_i$ | 0 (basal) or 10 (non-basal) | Maximum extinction rate due to the lack of prey |
| $M_i$ | 0.05 | Number of prey species giving the half-max extinction rate by lack of prey |
| $e_i$ | $\in \{0.1, 0.5, 0.9\}$ | Extinction rate because of disturbances |
| $h_j$ $(j = 1, 2, 3)$ | 1 | Hill coefficients to determine the forms of functions. |

Fig. S16 shows whether FCL in each definition correlated with three environmental drivers in the preliminary simulations or not. While resource availability did not correlate with FCL, disturbance and ecosystem size negatively and positively correlated with FCL, respectively, which justified our focus on these two environmental drivers in the main text. Correlations between FCL and resource availability slightly changed depending on disturbance and ecosystem size (Fig. S17). Although the effect of resource availability might also be contingent on the other environmental drivers, our model could not find enough strong correlations.

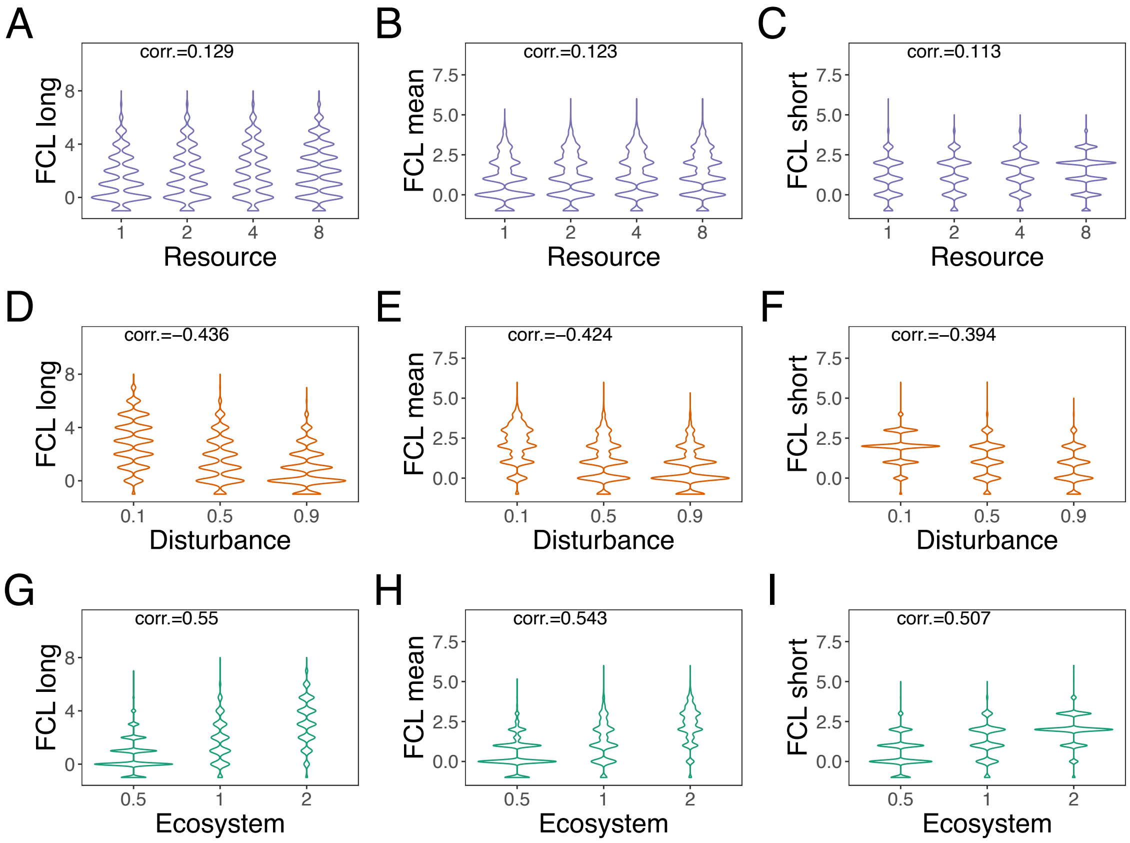


Figure S16: Correlations between FCL and environmental drivers

Each panel shows how distributions of FCL change over resource availability (A-C), disturbance (D-F), and ecosystem size (G-I). Spearman correlation coefficient appears on each panel. Corresponding p-values are omitted because they are small due to the large amount of data.

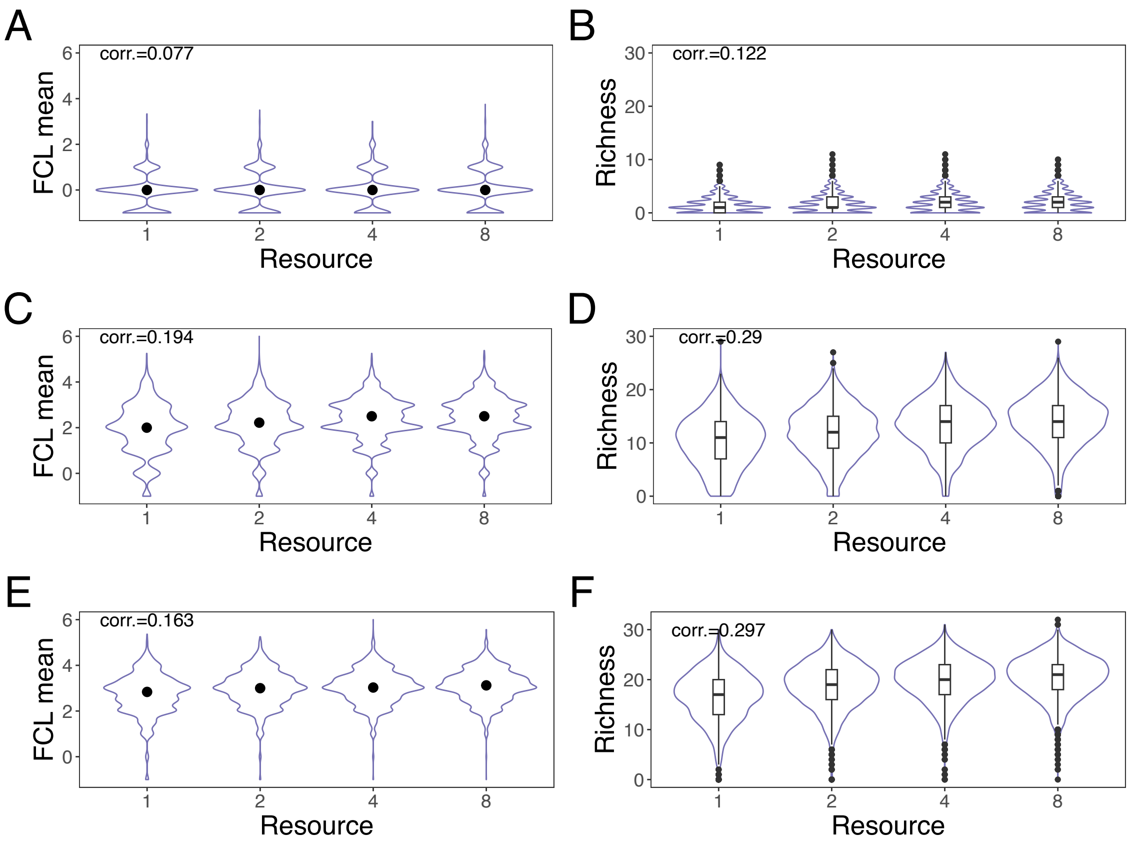


Figure S17: Potential effects of resource availability on FCL

Spearman correlations between resource availability and FCL mean (A, C, and E), or operational species richness (B, D, and F). In each row, disturbance and ecosystem size vary to change the median operational species richness. A and B: disturbance = 0.9 and ecosystem size = 0.5 generated low-richness communities (median operational species richness across resource availability is 2). Here, resource availability did not change either FCL or operational species richness. C and D: disturbance = 0.5 and ecosystem size = 2.0, resulting in median operational species richness = 13. In this case, resource availability positively correlated with FCL and operational species richness. E and F: disturbance = 0.1 and ecosystem size = 2.0. In this case, median operational species richness was so high (19), that the correlation between FCL and resource availability was weakened.

### SI 8 Supplimentary figures and tables

Table S3: Equations of the best models

| Data | Equations |
|---|---|
| Empirical FCL long | $\mathrm{FCL} = 3.94 \times \{1 - \exp(-0.09 \times \mathrm{richness})\} - 7.3 \times \mathrm{chain}^2 + 9.45 \times \mathrm{chain} - 1$ |
| Empirical FCL mean | $\mathrm{FCL} = 3.01 \times \{1 - \exp(-0.14 \times \mathrm{richness})\} - 2.82 \times \mathrm{chain}^2 + 4.92 \times \mathrm{chain} - 1$ |
| Empirical FCL short | $\mathrm{FCL} = 2.85 \times \{1 - \exp(-0.19 \times \mathrm{richness})\} + 2.12 \times \mathrm{chain} - 1$ |
| Simulated FCL long | $\mathrm{FCL} = 6.61 \times \{1 - \exp(-0.10 \times \mathrm{richness})\} - 0.41 \times \mathrm{chain}^2 + 1.74 \times \mathrm{chain} - 1$ |
| Simulated FCL mean | $\mathrm{FCL} = 3.88 \times \{1 - \exp(-0.20 \times \mathrm{richness})\} - 0.45 \times \mathrm{chain}^2 + 1.74 \times \mathrm{chain} - 1$ |
| Simulated FCL short | $\mathrm{FCL} = 3.03 \times \{1 - \exp(-0.30 \times \mathrm{richness})\} + 2.98 \times \{1 - \exp(-0.53 \times \mathrm{chain})\} - 1$ |

For simulations, we show only fixed effects.

Table S4: Linear quantile mixed model on median FCL mean

| Data | Variable | Coefficient | Standard error | P-value |
|---|---|---|---|---|
| Disturbance $\in [0.1, 0.3]$\|ecosystem size = 0.5 | Intercept | 2.50 | 0.10 | $< 10^{-5}$ |
| | Disturbance | $-3.75$ | 0.40 | $< 10^{-5}$ |
| Disturbance $\in [0.1, 0.3]$\|ecosystem size = 2.0 | Intercept | 3.36 | 0.13 | $< 10^{-5}$ |
| | Disturbance | $-0.65$ | 0.32 | $< 10^{-5}$ |
| Ecosystem size $\in [1.5, 2.0]$\|disturbance = 0.1 | Intercept | 2.80 | 0.13 | $1.06 \times 10^{-3}$ |
| | Ecosystem size | 0.31 | 0.09 | $6.78 \times 10^{-4}$ |
| Ecosystem size $\in [1.5, 2.0]$\|disturbance = 0.8 | Intercept | 1.14 | 0.31 | 0.016 |
| | Ecosystem size | 0.78 | 0.17 | $4.03 \times 10^{-5}$ |

We show only fixed effects.

Table S5: Linear quantile mixed model on median operational species richness

| Data | Variable | Coefficient | Standard error | P-value |
|---|---|---|---|---|
| Disturbance $\in [0.1, 0.3]$\|ecosystem size = 0.5 | Intercept | 8.00 | 0.10 | $< 10^{-5}$ |
| | Disturbance | $-10.00$ | 1.99 | $< 10^{-5}$ |
| Disturbance $\in [0.1, 0.3]$\|ecosystem size = 2.0 | Intercept | 22 | 0.62 | $< 10^{-5}$ |
| | Disturbance | $-16.67$ | 1.95 | $< 10^{-5}$ |
| Ecosystem size $\in [1.5, 2.0]$\|disturbance = 0.1 | Intercept | 8.50 | 1.65 | $< 10^{-5}$ |
| | Ecosystem size | 6.00 | 0.84 | $< 10^{-5}$ |
| Ecosystem size $\in [1.5, 2.0]$\|disturbance = 0.8 | Intercept | 0.36 | 1.29 | 0.78 |
| | Ecosystem size | 5.10 | 0.75 | $< \times 10^{-5}$ |

We show only fixed effects.

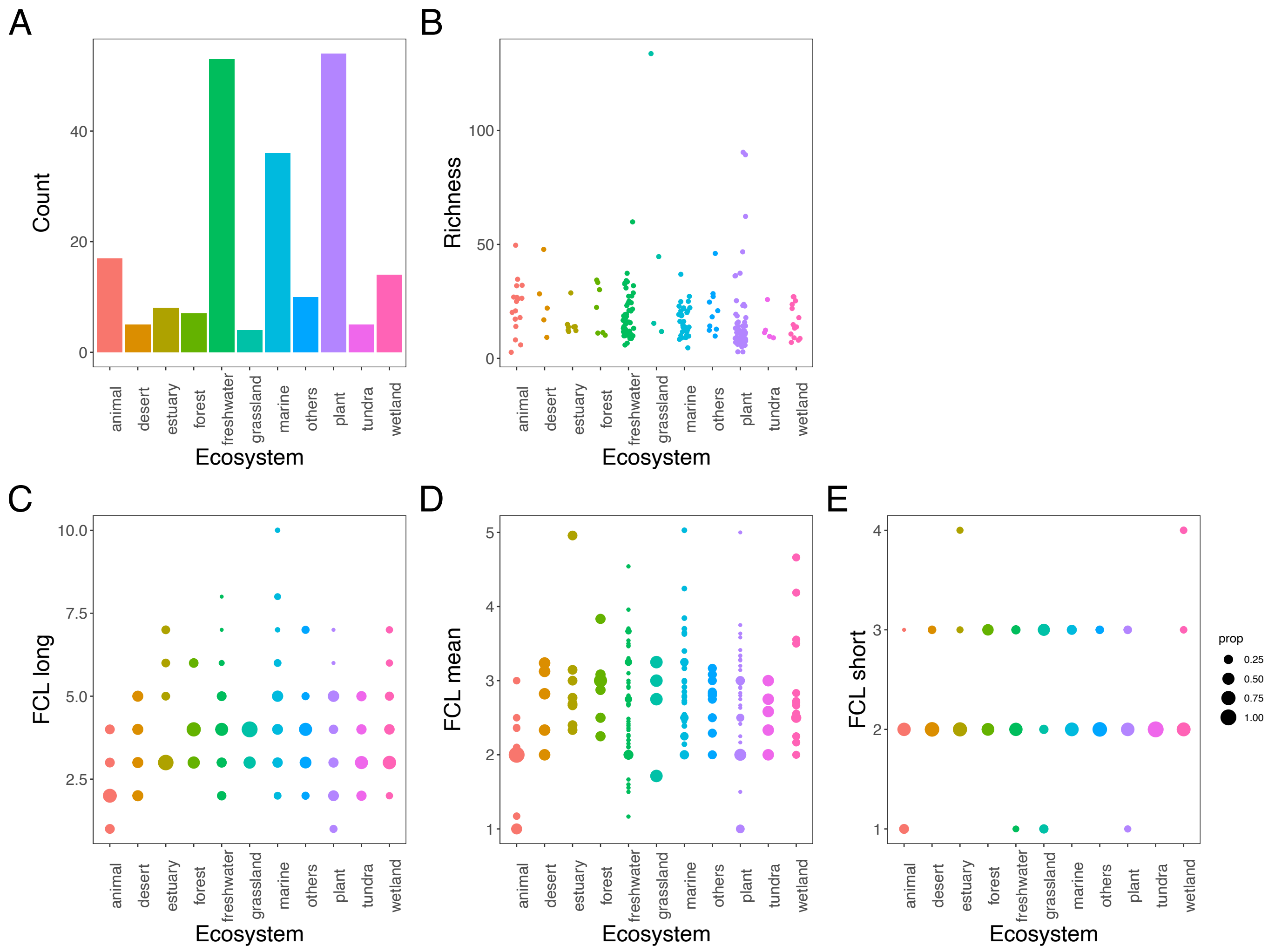


Figure S18: Ecosystem classes in the empirical data

A: The number of food webs in each ecosystem class: animal and plant represent food webs on certain animal bodies and on certain plant species, respectively. The class of "others" includes the following ecosystems: beach, island, mountain, old-field, and soil. B: operational species richness in each ecosystem class. C - E: FCL long(C), mean (D), and short (E) in each ecosystem class, respectively. The dots sizes follow the proportions of the amounts of data in each ecosystem.

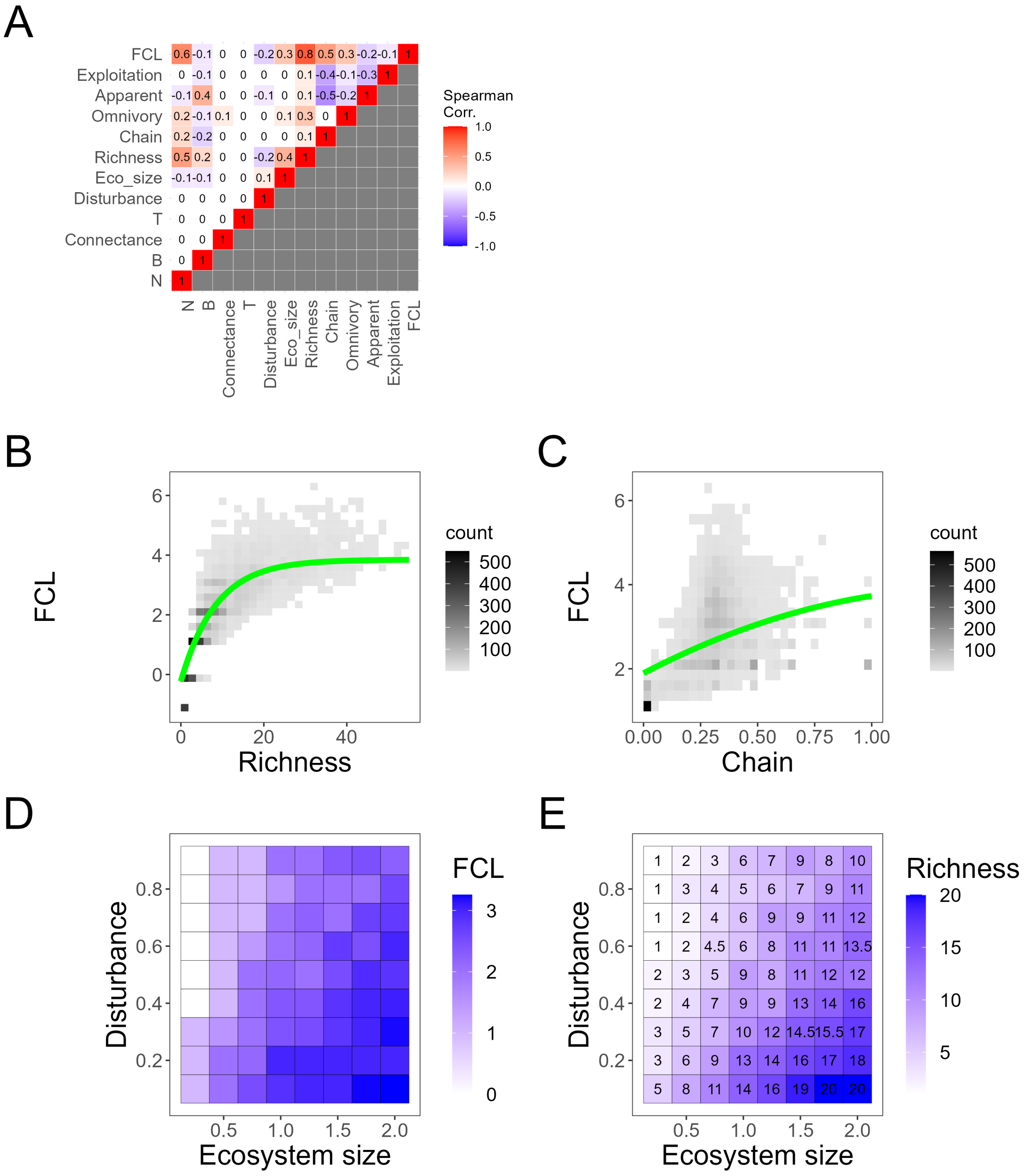


Figure S19: Sensitivity analysis in FCL mean

We performed the sensitivity analyses of our main results to parameters that affect food web structures: the maximum operational species richness $N$, the maximum number of basal species $B$, expected connectance in $N$-species food web, and the pattern of omnivory $T$. A: Spearman correlation coefficients are shown. FCL positively correlates with operational species richness and the fraction of chain motifs in realized food webs. B and C: Operational species richness and the fraction of chain motifs are fitted to the saturating and quadratic functions, respectively. The green curves represent $\text{FCL} = 4.08 \times (1 - \exp(-0.12 \times \text{richness})) - 0.98 \times \text{chain}^2 + 2.82 \times \text{chain} - 1$. On panel B, we fixed the fraction of chain motifs as its mean value, while we used mean richness on panel C. D and E: median FCL and operational species richness over two environmental drivers (disturbance and ecosystem size) are shown.

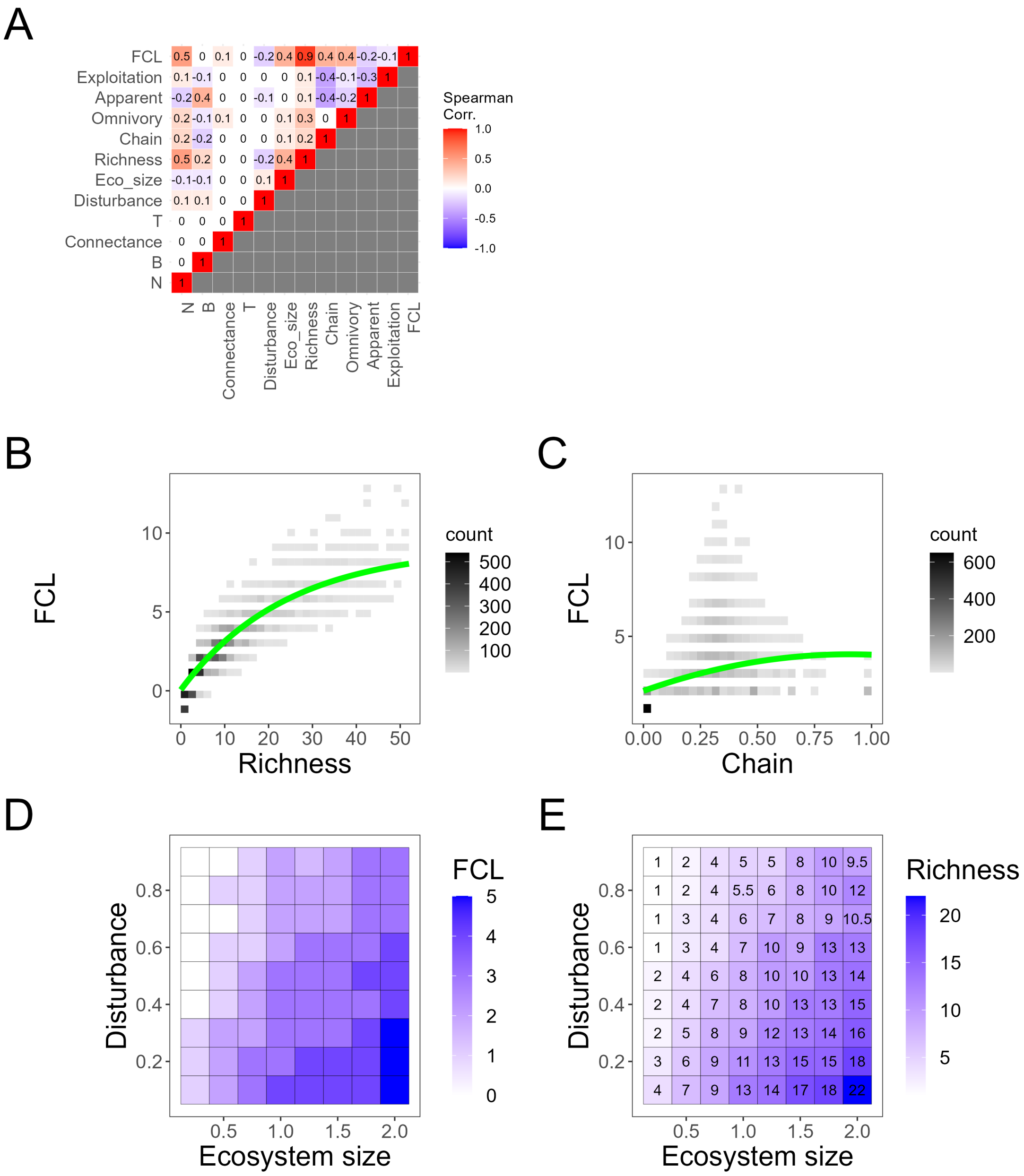


Figure S20: Sensitivity analysis in FCL long

Similar to Fig. S19, but this figure corresponds to FCL long. The green curves on panels B and C represent $\text{FCL} = 9.03 \times (1 - \exp(-0.04 \times \text{richness})) - 2.39 \times \text{chain}^2 + 4.31 \times \text{chain} - 1$.

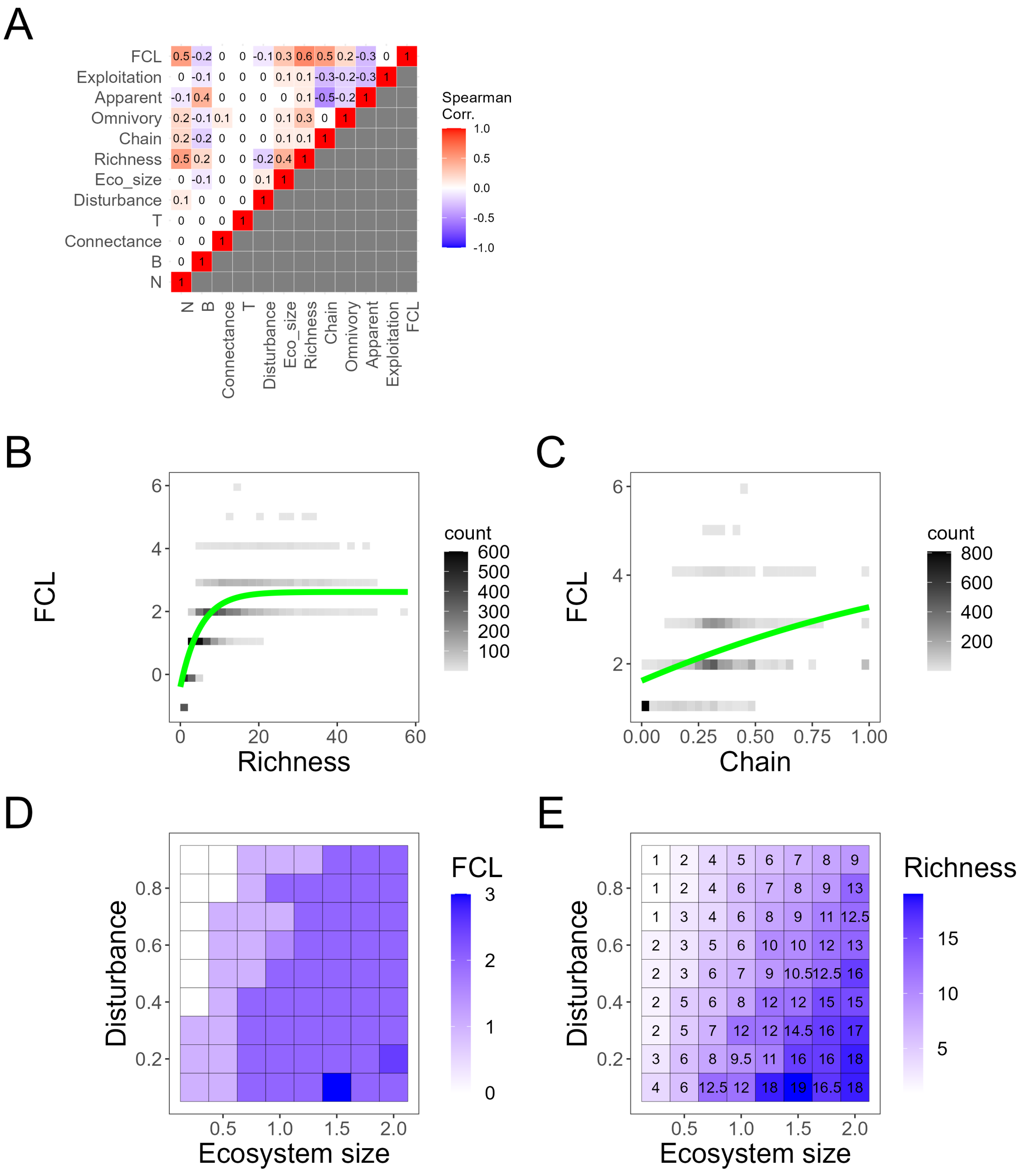

Figure S21: Sensitivity analysis in FCL short

Similar to Fig. S19, but this figure corresponds to FCL short. The green curves on panels B and C represent $FCL = 3.02 \times (1 - \exp(-0.20 \times \text{richness})) - 0.50 \times \text{chain}^2 + 2.16 \times \text{chain} - 1$.

Table S6: Model comparisons for empirical FCL long

| Model | ΔAIC |
|---|---|
| Saturating + Quadratic | 0.00 |
| Saturating + Saturating | 2.08 |
| Saturating + Linear | 14.40 |
| Linear + $\phi$ | 342.355 |
| Saturating + $\phi$ | 60.94 |
| Quadratic + $\phi$ | 196.32 |
| $\phi$ + Linear | 311.28 |
| $\phi$ + Saturating | 102.09 |
| $\phi$ + Quadratic | 342.46 |
| Linear + Linear | 102.28 |
| Linear + Saturating | 47.85 |
| Linear + Quadratic | 113.13 |
| Quadratic + Linear | 55.85 |
| Quadratic + Saturating | 17.92 |
| Quadratic + Quadratic | 26.75 |

In each row, the model represents the functional form of operational species richness (left) and the fraction of the chain motif (right). $\phi$ represents either operational species richness or the chain motif is not included in the model.

Table S7: Model comparisons for empirical FCL mean

| Model | ΔAIC |
|---|---|
| Saturating + Quadratic | 0.00 |
| Saturating + Saturating | 1.65 |
| Saturating + Linear | 11.73 |
| Linear + $\phi$ | 614.17 |
| Saturating + $\phi$ | 110.18 |
| Quadratic + $\phi$ | 438.35 |
| $\phi$ + Linear | 609.00 |
| $\phi$ + Saturating | 240.69 |
| $\phi$ + Quadratic | 486.60 |
| Linear + Linear | 351.63 |
| Linear + Saturating | 171.14 |
| Linear + Quadratic | 255.62 |
| Quadratic + Linear | 164.63 |
| Quadratic + Saturating | 82.32 |
| Quadratic + QUadratic | 89.35 |

In each row, the model represents the functional form of operational species richness (left) and the fraction of the chain motif (right). $\phi$ represents either operational species richness or the chain motif is not included in the model.

Table S8: Model comparisons for empirical FCL short

| Model | ΔAIC |
|---|---|
| Saturating + Linear | 0.00 |
| Saturating + Quadratic | 1.34 |
| Saturating + Saturating | 1.40 |
| Linear + $\phi$ | 586.39 |
| Saturating + $\phi$ | 65.68 |
| Quadratic + $\phi$ | 407.15 |
| $\phi$ + Linear | 596.21 |
| $\phi$ + Saturating | 251.17 |
| $\phi$ + Quadratic | 437.51 |
| Linear + Linear | 1365.91 |
| Linear + Saturating | 183.29 |
| Linear + Quadratic | 234.91 |
| Quadratic + Linear | 155.35 |
| Quadratic + Saturating | 100.03 |
| Quadratic + QUadratic | 108.9 |

In each row, the model represents the functional form of operational species richness (left) and the fraction of the chain motif (right). $\phi$ represents either operational species richness or the chain motif is not included in the model.

Table S9: Model comparisons for simulated FCL long

| Model | ΔAIC |
|---|---|
| Saturating + Quadratic | 0.00 |
| Saturating + Saturating | 50 |
| Saturating + Linear | 19430 |
| Linear + $\phi$ | 44905 |
| Saturating + $\phi$ | 6991 |
| Quadratic + $\phi$ | 12486 |
| $\phi$ + Linear | 105663 |
| $\phi$ + Saturating | 67799 |
| $\phi$ + Quadratic | 72435 |
| Linear + Linear | 25893 |
| Linear + Saturating | 23119 |
| Linear + Quadratic | 23102 |
| Quadratic + Linear | 4270 |
| Quadratic + Saturating | 4280 |
| Quadratic + QUadratic | 4236 |

In each row, the model represents the functional form of operational species richness (left) and the fraction of the chain motif (right). $\phi$ represents either operational species richness or the chain motif is not included in the model.

Table S10: Model comparisons for simulated FCL mean

| Model | ΔAIC |
|---|---|
| Saturating + Quadratic | 0.00 |
| Saturating + Linear | 148 |
| Saturating + Satuating | 1582 |
| Linear + $\phi$ | 82123 |
| Saturating + $\phi$ | 12357 |
| Quadratic + $\phi$ | 30528 |
| $\phi$ + Linear | 114416 |
| $\phi$ + Saturating | 77980 |
| $\phi$ + Quadratic | 83838 |
| Linear + Linear | 57900 |
| Linear + Saturating | 54194 |
| Linear + Quadratic | 54294 |
| Quadratic + Linear | 18503 |
| Quadratic + Saturating | 18362 |
| Quadratic + QUadratic | 18268 |

In each row, the model represents the functional form of operational species richness (left) and the fraction of the chain motif (right). $\phi$ represents either operational species richness or the chain motif is not included in the model.

Table S11: Model comparisons for simulated FCL short

| Model | ΔAIC |
|---|---|
| Saturating + Saturating | 0.00 |
| Saturating + Quadratic | 663 |
| Saturating + Linear | 1460 |
| Linear + $\phi$ | 79060 |
| Saturating + $\phi$ | 12095 |
| Quadratic + $\phi$ | 31753 |
| $\phi$ + Linear | 96344 |
| $\phi$ + Saturating | 64524 |
| $\phi$ + Quadratic | 70139 |
| Linear + Linear | 55177 |
| Linear + Saturating | 51981 |
| Linear + Quadratic | 52219 |
| Quadratic + Linear | 21799 |
| Quadratic + Saturating | 21356 |
| Quadratic + QUadratic | 21095 |

In each row, the model represents the functional form of operational species richness (left) and the fraction of the chain motif (right). $\phi$ represents either operational species richness or the chain motif is not included in the model.

Table S12: Model comparisons for simulated FCL long with the alternative implementation of ecosystem size

| Model | ΔAIC |
|---|---|
| Saturating + Quadratic | 0.00 |
| Saturating + Saturating | 34854 |
| Saturating + Linear | 2985 |
| Linear + $\phi$ | 46180 |
| Saturating + $\phi$ | 1128 |
| Quadratic + $\phi$ | 16010 |
| $\phi$ + Linear | 117656 |
| $\phi$ + Saturating | 78348 |
| $\phi$ + Quadratic | 85287 |
| Linear + Linear | 25948 |
| Linear + Saturating | 24593 |
| Linear + Quadratic | 524422 |
| Quadratic + Linear | 4400 |
| Quadratic + Saturating | 4369 |
| Quadratic + QUadratic | 4306 |

In each row, the model represents the functional form of operational species richness (left) and the fraction of the chain motif (right). $\phi$ represents either operational species richness or the chain motif is not included in the model.

Table S13: Model comparisons for simulated FCL mean with the alternative implementation of ecosystem size

| Model | ΔAIC |
|---|---|
| Saturating + Quadratic | 0.00 |
| Saturating + Saturating | 236 |
| Saturating + Linear | 771 |
| Linear + $\phi$ | 86279 |
| Saturating + $\phi$ | 20088 |
| Quadratic + $\phi$ | 35067 |
| $\phi$ + Linear | 132182 |
| $\phi$ + Saturating | 94103 |
| $\phi$ + Quadratic | 101227 |
| Linear + Linear | 60688 |
| Linear + Saturating | 58721 |
| Linear + Quadratic | 58403 |
| Quadratic + Linear | 17614 |
| Quadratic + Saturating | 17574 |
| Quadratic + QUadratic | 17508 |

In each row, the model represents the functional form of operational species richness (left) and the fraction of the chain motif (right). $\phi$ represents either operational species richness or the chain motif is not included in the model.

Table S14: Model comparisons for simulated FCL short with the alternative implementation of ecosystem size

| Model | ΔAIC |
|---|---|
| Saturating + Saturating | 0.00 |
| Saturating + Quadratic | 500 |
| Saturating + Linear | 518 |
| Linear + $\phi$ | 86832 |
| Saturating + $\phi$ | 16740 |
| Quadratic + $\phi$ | 36271 |
| $\phi$ + Linear | 114657 |
| $\phi$ + Saturating | 79759 |
| $\phi$ + Quadratic | 86609 |
| Linear + Linear | 60388 |
| Linear + Saturating | 58435 |
| Linear + Quadratic | 58468 |
| Quadratic + Linear | 21130 |
| Quadratic + Saturating | 20736 |
| Quadratic + QUadratic | 20499 |

In each row, the model represents the functional form of operational species richness (left) and the fraction of the chain motif (right). $\phi$ represents either operational species richness or the chain motif is not included in the model.

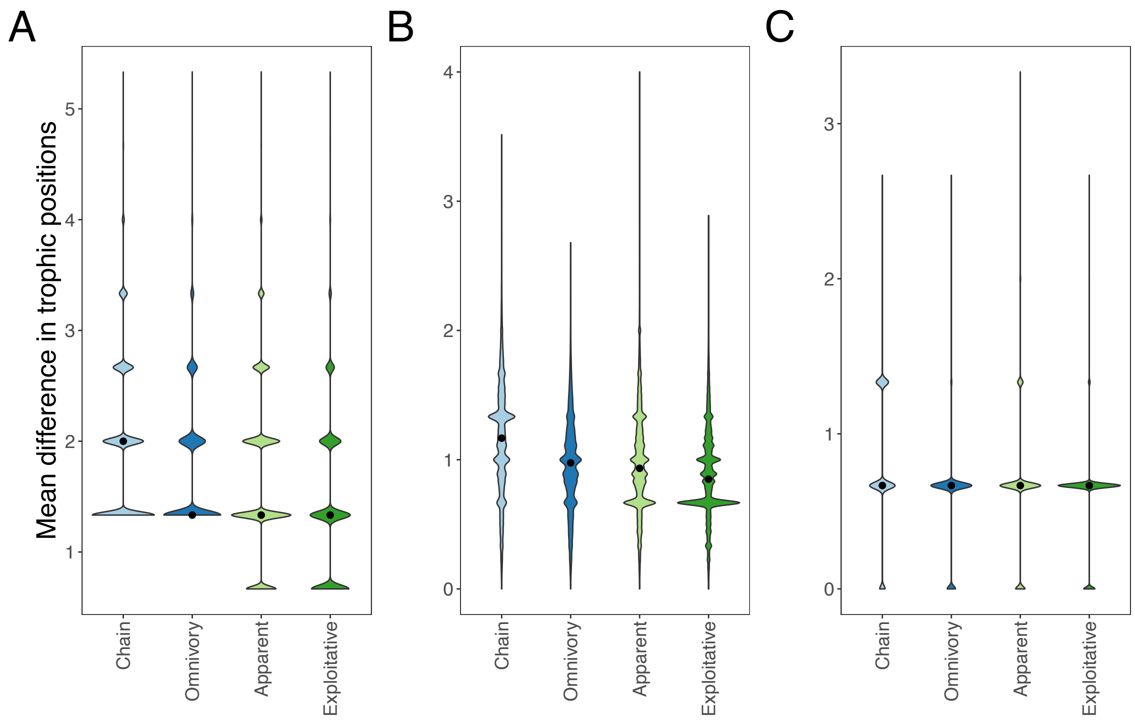


Figure S22: Mean difference in trophic positions within food web motifs

Distributions of mean difference in trophic levels within each food web motif in our simulations are shown. A: trophic positions are given by one plus the longest-path length from basal species, B: trophic positions are given by one plus mean trophic positions of prey species, and C: trophic positions are given by one plus the shortest-path length from basal species. The black dots represent median values. In all cases, the chain motifs have larger differences than other motifs (Wilcoxon rank-sum test, p-values $< 0.001$). See Table S15 for effect sizes (Cliff's delta).

Table S15: Cliff's delta of mean trophic position differences

| Pairs | FCL long | FCL mean | FCL short |
|---|---|---|---|
| Chain − Omnivory | 0.140 | 0.280 | 0.264 |
| Chain − Apparent | 0.226 | 0.258 | 0.186 |
| Chain − Exploitative | 0.414 | 0.380 | 0.244 |
